# Central opioid receptors mediate morphine-induced itch and chronic itch

**DOI:** 10.1101/2020.05.31.126805

**Authors:** Zilong Wang, Changyu Jiang, Hongyu Yao, Ouyang Chen, Sreya Rahman, Yun Gu, Yul Huh, Ru-Rong Ji

**Affiliations:** Center for Translational Pain Medicine, Department of Anesthesiology, Duke University Medical Center, Durham, North Carolina, 27710; Department of Cell Biology, Duke University Medical Center, Durham, North Carolina, 27710; Department of Neurobiology, Duke University Medical Center, Durham, North Carolina, 27710

**Author notes:** Correspondence should be addressed to: Ru-Rong Ji, PhD, Department of Anesthesiology, Duke University Medical Center, Durham, North Carolina, 27710., Or Zilong Wang, Department of Anesthesiology, Duke University Medical Center, Durham, North Carolina, 27710.

**Keywords:** allergic contact dermatitis, inhibitory interneurons, itch, lymphoma, opioid, spinal cord

## Abstract

Opioids, such as morphine are mainstay treatments for clinical pain conditions. Itch is a common side effect of opioids, particularly as a result of epidural or intrathecal (i.t.) administration. Recent progress has advanced our understanding of itch circuits in the spinal cord. However, the mechanisms underlying opioid-induced itch are not fully understood, although an interaction between µ-opioid receptor (MOR) and gastrin-releasing peptide receptor (GRPR) in spinal GRPR-expressing neurons has been implicated. In this study we investigated the cellular mechanisms of intrathecal (i.t.) opioid-induced itch by conditional deletion of MOR-encoding *Oprm1* in distinct populations of interneurons and sensory neurons. We found that i.t. injection of the MOR agonists morphine or DAMGO elicited dose-dependent scratching, but this pruritus was totally abolished in mice with a specific *Oprm1* deletion in Vgat^+^ neurons (*Oprm1*-*Vgat*). Loss of MOR in somatostatin^+^ interneurons and TRPV1^+^ sensory neurons did not affect morphine-induced itch but impaired morphine-induced antinociception. *In situ* hybridization revealed *Oprm1* expression in 30% of inhibitory and 20% of excitatory interneurons in the spinal dorsal horn. Whole-cell recordings from spinal cord slices showed that DAMGO induced outward currents in 9 out of 19 Vgat^+^ interneurons examined. Morphine also inhibited action potentials in Vgat^+^ interneurons and suppressed evoked IPSCs in postsynaptic Vgat^-^ excitatory neurons, suggesting a mechanism of disinhibition by MOR agonists. Notably, morphine-elicited itch was suppressed by i.t. administration of NPY and abolished by spinal ablation of GRPR^+^ neurons, whereas i.t. GRP-induced itch response remained intact in mice lacking *Oprm1*-*Vgat*. Additionally, chronic itch from DNFB-induced allergic contact dermatitis was decreased by *Oprm1*-*Vgat* deletion. Finally, naloxone, but not peripherally restricted naloxone methiodide, inhibited chronic itch in the DNFB model and the cutaneous T-cell lymphoma (CTCL) model, indicating a contribution of central MOR signaling to chronic itch. Our findings demonstrate that i.t. morphine elicits itch via acting on MOR on spinal inhibitory interneurons, leading to disinhibition of the spinal itch circuit. Our data also suggest that chronic itch could be effectively treated with CNS-targeted naloxone.

## Introduction

Opioids are mainstay pain treatments in clinical medicine. Most opioid analgesics produce antinociception via the µ-opioid receptor (MOR) which is expressed in both the peripheral and central nervous systems (Ji *et al*., 1995; Matthes *et al*., 1996; Corder *et al*., 2018). MOR mediates beneficial effects of opioid analgesics such as antinociception as well as unwanted side effects such as hyperalgesia, opioid induced constipation, and withdrawal responses. Itch is a notable side effect of opioids, particularly following epidural or intrathecal administration. The incidence of pruritus in patients treated systemically with opioids is about 2-10%, whereas the incidence of pruritus increases to 30-60% of patients receiving intrathecal opioid treatment. Pregnant women have been observed to be more susceptible to pruritus after spinal opioid administration, with an incidence of 60-100% (Reich and Szepietowski, 2010; Kumar and Singh, 2013).

Recent progress has advanced our understanding of the mechanisms of itch (LaMotte *et al*., 2014; Ji, 2018; Cevikbas and Lerner, 2019). Distinct populations of primary pruriceptors have been demonstrated to sense itch signals (Liu *et al*., 2009; Han *et al*., 2013; Mishra and Hoon, 2013; Qu *et al*., 2014; Pan *et al*., 2019). Several neurotransmitters and neuromodulators, as well as the involved neurocircuits in itch transmission have been identified (Sun and Chen, 2007; Sun *et al*., 2009; Carstens *et al*., 2010; Mishra and Hoon, 2013; Kardon *et al*., 2014; LaMotte *et al*., 2014; Huang *et al*., 2018). Recently, two populations of inhibitory interneurons expressing Bhlhb5 and NPY in the spinal dorsal horn (SDH) have been implicated in regulating chemical and mechanical itch, respectively (Ross *et al*., 2010; Bourane *et al*., 2015; Acton *et al*., 2019; Pan *et al*., 2019).

Patients with chronic itch commonly experience high sensitivity to pruritogens, mechanically evoked itch sensations, and spontaneous itch (Ikoma *et al*., 2006; LaMotte *et al*., 2014). Opioid receptor antagonists (e.g., naloxone, naltrexone, and nalbuphine) have shown to be effective for chronic itch following dermatitis, uremic pruritus, and anti-PD1 immunotherapy induced itch (Brune *et al*., 2004; Kwatra *et al*., 2018; Reszke and Szepietowski, 2018; Serrano *et* al., 2018; Kremer, 2019; Singh *et al*., 2019). However, we know little about the molecular mechanisms underlying opioid-induced itch beyond a demonstrated interaction between µ-opioid receptor isoform 1D (MOR1D) and gastrin-releasing peptide receptor (GRPR) (Liu *et al*., 2011). To tackle this problem, we conditionally knocked out MOR in different populations of itch-modulating neurons, including nociceptive sensory neurons expressing transient receptor potential ion channel subtype V1 (TRPV1), inhibitory neurons (Vgat^+^), and excitatory interneurons in the spinal cord dorsal horn (SDH) expressing somatostatin (SST^+^). We demonstrated that MOR in inhibitory interneurons, but not in excitatory interneurons of the SDH, mediates itch following i.t. µ-opioid treatment. Furthermore, MOR expression in spinal inhibitory interneurons is essential for driving dermatitis-associated chronic itch.

## Materials and methods

### Reagents

Morphine sulfate was obtained from WEST-WARD pharmaceuticals. Naloxone methiodide (Catalog: N192), naloxone (Catalog: 1453005), DAMGO (Catalog: E7384) GRP (Catalog: G8022), histamine (Catalog:H7125), chloroquine (Catalog: 1118000), and 1-Fluoro-2,4-dinitrobenzene (Catalog: D1529) were obtained from Sigma Aldrich. CTOP (Catalog: ab120417) was purchased from Abcam. RTX (Catalog: R8756). Bombesin-saporin (Catalog: IT-40) and blank-saporin (Catalog: IT-21) were purchased from Advanced targeting systems.

### Animals

*Oprm1*^fl/fl^ (stock No: 030074), *Vgat*-ires-Cre (stock No: 016962), *Sst*-ires-Cre (stock No:013044), *Trpv1*-Cre (stock No:017769), Ai32 (stock No:024109), C57BL/6J (stock No:000664), NOD. CB-17-Prkdcscid mice (stock No:001303), and Ai9 tdTomato (stock No:007909) mice were purchased from the Jackson Laboratory and maintained at the Duke University Animal Facility. Young mice (1-2 months of age for both sexes) were used for electrophysiological studies of spinal cord tissue. Adult male and female mice (2-4 months), including knockout mice and corresponding wild-type control mice, as well as CD1 mice (Fig. 2D) were used for behavioral and pharmacological studies. Mice were group-housed on a 12-hour light/12-hour dark cycle at 22±1 °C with free access to food and water. All mice were randomized when assorted for animal experiments. Sample sizes were estimated based on our previous studies for similar types of behavioral, biochemical, and electrophysiological assays and analyses (Supplementary Table 1) (Chen *et al*., 2014; Han *et al*., 2018; Wang *et al*., 2020). Two to five mice were housed per cage. The animal studies were approved by Institutional Animal Care and Use Committee (IACUC) of Duke University. Animal experiments were conducted in accordance with the National Institutes of Health Guide for the Care and Use of Laboratory Animals and ARRIVE guidelines.

**Figure 1.**
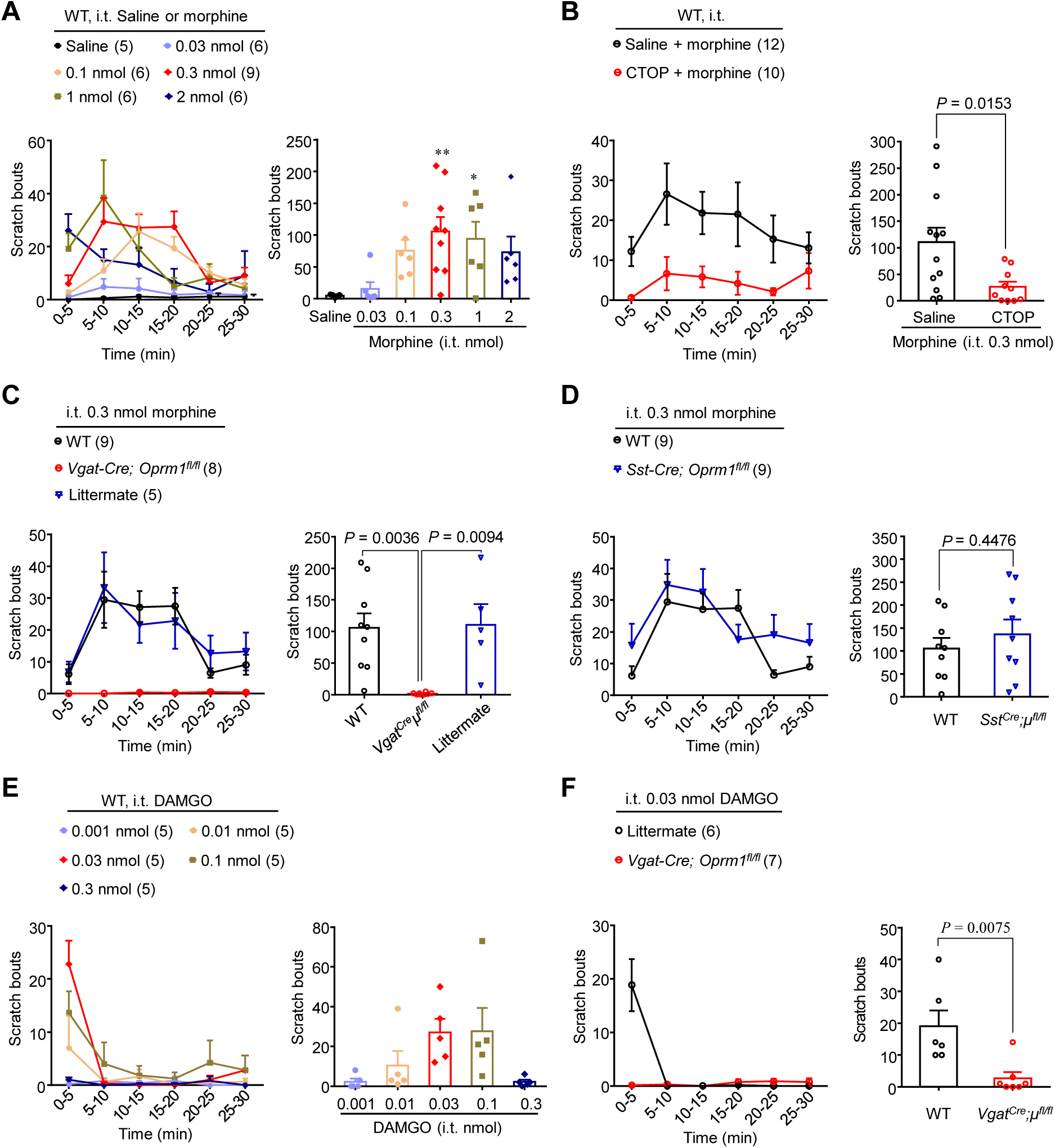
Intrathecal morphine-induced itch requires MOR in spinal inhibitory interneurons. **(A)** Time course (left) and total scratch bouts (right) within 30 min after intrathecal injection of different doses of morphine in WT mice. \**P* = 0.0392, \*\**P* = 0.0086, One-way ANOVA test, followed by Dunnett’s post hoc test. **(B)** Pretreatments with MOR selective antagonist CTOP (0.15 pmol, i.t.) blocked the morphine (0.03 nmol, i.t.) induced itch response. *P* = 0.0153, two-tailed Student’s t-test. **(C)** Intrathecal morphine-induced itch was totally abolished in *Vgat-Cre; Oprm1*^*fl/fl*^ mice. *P* = 0.0036 vs. WT mice, *P* = 0.0094 vs. Cre negative littermates, two-tailed Student’s t-test. **(D)** Intrathecal morphine-induced itch was not changed in *Sst-Cre; Oprm1*^*fl/fl*^ mice. *P* = 0.4476, two-tailed Student’s t-test. **(E)** Time course (left) and total scratch bouts (right) within 30 min after intrathecal injection of different doses of MOR selective agonist DAMGO in WT mice. **(F)** Intrathecal 0.03 nmol DAMGO induced itch was totally abolished in *Vgat-Cre; Oprm1*^*fl/fl*^ mice. *P* = 0.0075 vs. Cre negative littermates, two-tailed Student’s t-test. Data are mean ± SEM. Sample sizes are indicated in the figures.

**Figure 2.**
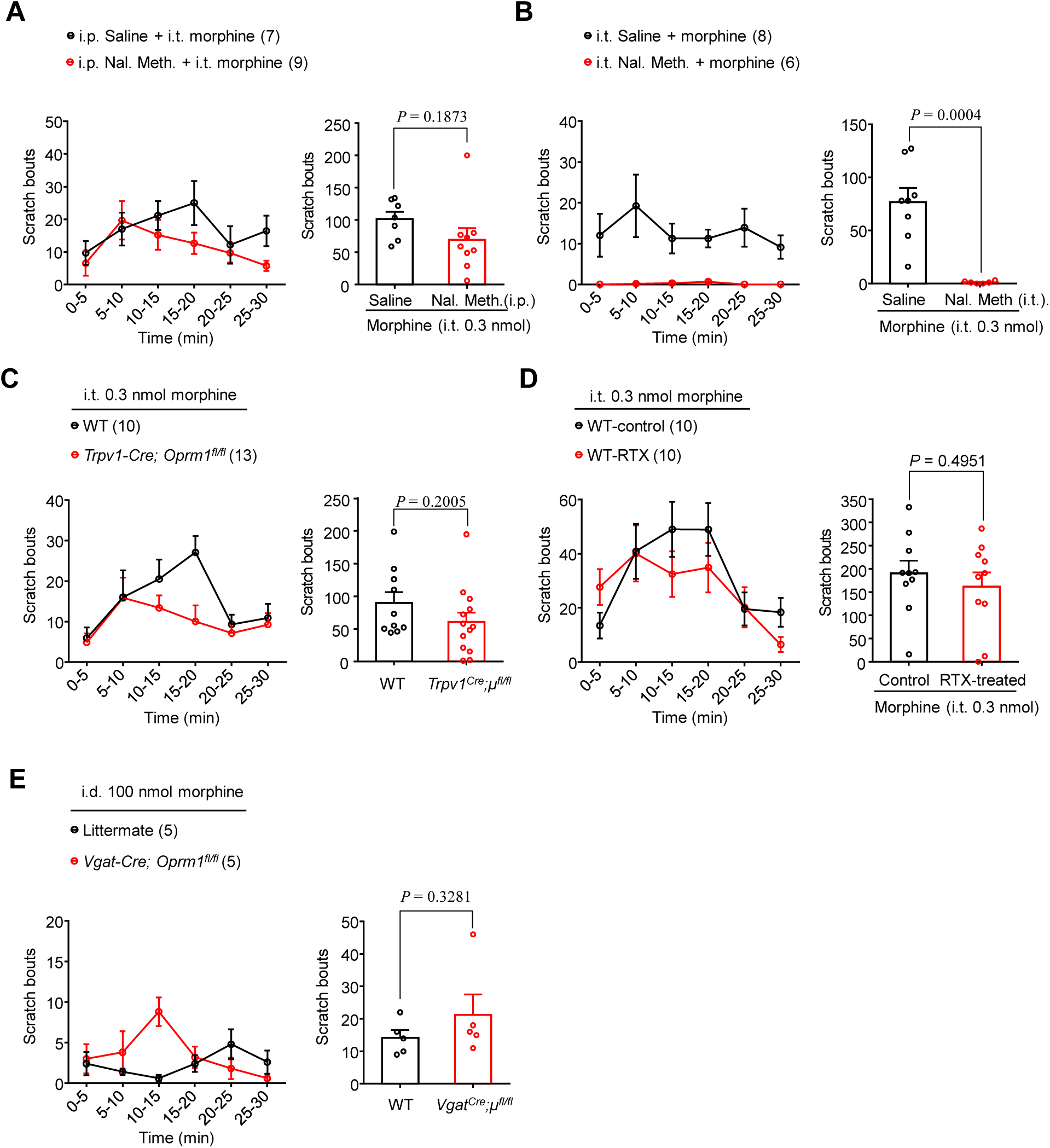
Intrathecal morphine-induced itch does not require peripheral MOR. **(A)** Intraperitoneal pretreatment with peripherally restricted MOR antagonist naloxone methiodide (10 mg/kg) did not affect itch induced by i.t. morphine (0.3 nmol). *P* = 0.1873, two-tailed Student’s t-test. **(B)** Pretreatment with naloxone methiodide (5 µg, i.t.) totally blocked intrathecal morphine-induced pruritus. *P* = 0.0004, two-tailed Student’s t-test. **(C)** Intrathecal morphine-induced pruritus was not altered in *Trpv1-Cre; Oprm1*^*fl/fl*^ mice. *P* = 0.2005, two-tailed Student’s t-test. **(D)** RTX ablation of TRPV1^+^ sensory neurons did not change intrathecal morphine-induced itch response. *P* = 0.4951, two-tailed Student’s t-test. **(E)** Intradermal injection of morphine induced a very mild scratch response that was not affected in *Vgat-cre; Oprm1*^*fl/fl*^ mice. *P* = 0.3281, two-tailed Student’s t-test. Data are mean ± SEM. Sample sizes are indicated in the figures.

### Drug injection

For intrathecal injections, lumbar puncture was made by a Hamilton microsyringe (Hamilton) fitted with a 30-G needle between the L5 and L6 spinal levels to deliver reagents (5 µl) into cerebral spinal fluid. For intradermal administration, a volume of 50 µl of pruritogen was injected into the skin of the nape with a 30-G needle.

### Behavioral assessment for itch response

Mice were shaved on the nape or back under light anesthesia with isoflurane. Before experiments, mice were habituated in small plastic chambers (14 × 18 × 12 cm) for two days. Room temperature and humidity levels remained constant and stable for all experiments. Mice were given intrathecal injections of morphine, DAMGO, or GRP, or intradermal injections of morphine, histamine, or chloroquine (CQ). After injection, scratching behavior was video recorded for 30 min using a Sony HDR-CX610 camera. The video was subsequently played back and the number of scratches by each mouse were quantified in a blinded manner. A scratch was counted when a mouse lifted its hind paw to scratch the injected region (for intradermal injection) or the back (for intrathecal injection).

### DNFB-induced allergic contact dermatitis in neck skin

The allergic contact dermatitis (ACD) model of chronic itch was generated by applying the hapten 1-fluoro-2, 4-dinitrobenzene (DNFB) onto the back skin as previously described (Liu *et al*., 2016). DNFB was dissolved in a mixture of acetone and olive oil in a 4:1 ratio. The surface of the abdomen and the nape of neck of each mouse were shaved 1 day before sensitization. On day 0, mice were sensitized with 50 μl of 0.5% DNFB solution by topical application to a 2 cm^2^ area of shaved abdomen skin. On days 5, 7, 9, and 11 mice were challenged with 30 μl 0.25% DNFB solution painted onto the nape of neck, with 1-hour videos taken on days 6, 8, 10 and 12. Spontaneous scratching was observed for each 1 hour recording.

### Cutaneous T cell lymphoma (CTCL) model

The CTCL model was established in a similar method as our previous study (Han *et al*., 2018). Briefly, a CD4^+^ MyLa cell line was purchased from Sigma (Ca#95051032). The cell line was established from a plaque biopsy of an 82-year old male with mycosis fungoides stage II by inclusion of IL-2 and IL-4 in the culture medium. CTCL was generated by intradermal injection of CD4^+^ Myla cells (1× 10^5^ cells/μl, 100 μl) on the nape of the neck in immune-deficient mice (NOD.CB17-Prkdcscid, 8–10 weeks old, female and male). Itch behavior was tested 20 – 30 days after MyLa cell inoculation and the videos were analyzed (1 hour per analysis).

### Tail-flick test

All animals were habituated to the testing environment for at least 2 days before baseline testing. Tail-flick testing was performed as described in a previous report (Wang *et al*., 2020). Briefly, mice were gently held by hand with a terry glove with tail exposed. The distal 3 cm end of the tail was immersed into a 50°C hot water bath. The tail-flick latency was measured as the time required for the mouse to flick or remove its tail from the water, with a maximum cut-off value of 15 seconds to prevent thermal injury. Tail-flick latency was determined both before and after drug injection. Data are expressed as the maximum possible effect (MPE) where MPE (%) = 100 × [(postdrug response – baseline response)/(cutoff response − baseline response)]. The MPE (%) value from each animal was converted to an area under the curve (AUC) value.

### Hot-plate test

All animals were habituated to the testing environment for at least 2 days before baseline testing. Hot-plate testing was conducted following tail-flick testing. Mice were placed on the hot plate apparatus set at 53°C, and the reaction time was scored when the animal began to exhibit signs of pain avoidance such as jumping or paw licking. A maximum cut-off value of 40 seconds was set to avoid thermal injury.

### *In situ* hybridization

Animals were deeply anesthetized with isoflurane and transcardiac perfusion was performed with PBS, followed by fixation with 4% paraformaldehyde. After perfusion and fixation, spinal cords were removed and post-fixed in 4% paraformaldehyde overnight at 4 °C. The tissues were then cryopreserved in 20% sucrose in PBS for 1 day followed by 30% sucrose in PBS for 1 day. Spinal cord sections (20 µm) were cut using a cryostat. *In situ* hybridization (ISH) was performed using the RNAscope system (Advanced Cell Diagnostics) following the manufacturer’s protocol and our previous reports (Chen *et al*., 2017; Wang *et al*., 2020). Pretreatment consisted of dehydration, followed by incubation with hydrogen peroxide and protease IV at room temperature. The Multiplex Fluorescent Kit v2 protocol was followed using commercial probes for MOR (Mm-*Oprm1*-C3, # 315841-C3), VGLUT2 (Mm-*Slc17a6*, #319171-C2), NPY (Mm-*Npy*, #313321-C2), NPY1R (Mm-*Npy1r*, #427021), PDYN (Mm-*Pdyn*, #318771), GRP (Mm-*Grp*, #317961-C2), and GRPR (Mm-*Grpr*, #317871). ISH images were captured by Zeiss 880 inverted confocal microscopy. For quantification purposes, all images acquired with the same settings, 2 to 3 sections from each animal were selected, and a total of four animals were included for data analysis. Visualized cells with more than 5 puncta per cell were classified as positive neurons.

### Spinal cord slice preparation and patch-clamp recordings

Mice were anesthetized with urethane (1.5-2.0 g/kg, i.p.), the lumbosacral spinal cord was quickly dissected, and the tissue was placed in ice-cold dissection solution (mM: Sucrose 240, NaHCO_3_ 25, KCl 2.5, NaH_2_PO_4_ 1.25, CaCl_2_ 0.5 and MgCl_2_ 3.5), equilibrated with 95% O_2_ and 5% CO_2_. Mice were sacrificed by decapitation following spinal extraction under anesthesia. Transverse spinal slices (300-400 μm) were cut using a vibrating microslicer (VT1200s Leica). The slices were incubated at 32°C for 1 hour in ACSF (mM: NaCl 126, KCl 3, MgCl_2_ 1.3, CaCl2 2.5, NaHCO_3_ 26, NaH2PO4 1.25 and glucose 11), equilibrated with 95% O_2_ and 5% CO_2_. The slices were then placed in a recording chamber and perfused at a flow rate of 2-4 ml/min with ACSF which was saturated with 95% O_2_ and 5% CO_2_ and maintained at room temperature (Jiang *et al*., 2014). Lamina II neurons in lumbar segments were identified as a translucent band under a microscope (BX51WIF; Olympus) with light transmitted from below. Vgat^+^ neurons were identified by observed fluorescence.

Whole-cell voltage-clamp recordings were made from lamina II neurons by using patch-pipettes fabricated from thin-walled, fiber-filled capillaries. The patch-pipette solution used to record opioid-induced currents or action potentials contained: K-gluconate 135, KCl 5, CaCl_2_ 0.5, MgCl_2_ 2, EGTA 5, HEPES 5, Mg-ATP 5 (pH=7.3). DAMGO-induced currents were recorded at a holding potential of -70 mV in voltage clamp mode. Action potentials were recorded in current clamp mode. To record light-evoked IPSCs, the patch-pipette solution contained (Gao *et al*.): Cs_2_SO_4_ 110, CaCl_2_ 0.5, MgCl_2_ 2, EGTA 5, HEPES 5, Mg-ATP 5, tetraethylammonium (TEA)-Cl 5 (pH=7.3). For optogenetic activation of inhibitory interneurons, *Vgat*-Cre; Ai32 mice were used. Blue light illumination (0.1 s) was delivered through a 40X water-immersion microscope (BX51WIF; Olympus) objective to initiate a light-evoked response. The light-evoked IPSCs were recorded in the presence of AP-V (50 μM) and CNQX (2 μM) at a holding potential of 0 mV in voltage clamp mode. The patch-pipettes had a resistance of 8-10 MΩ. Signals were acquired using an Axopatch 700B amplifier. The data were analyzed with pCLAMP 10.3 software. Currents were measured by Clampfit. Numerical data are given as the mean ± SEM. In all cases, *n* refers to the number of the neurons studied. All drugs were bath applied by gravity perfusion via a three-way stopcock without any change to the perfusion rate.

### Statistical analyses

All data are expressed as mean ± SEM as indicated in the figure legends. Statistical analyses were completed with Prism GraphPad 6.1. Data were analyzed using two-tailed student’s t-test (two groups), One-Way ANOVA, and Two-Way ANOVA. The criterion for statistical significance was *P* < 0.05.

### Data availability

Original data are available upon request.

## Results

### Intrathecal opioids induced acute itch via MOR expressed by inhibitory interneurons

We first tested intrathecally injected morphine-induced itch response in WT mice. Intrathecal morphine elicited dose-dependent scratch behavior in WT mice (Fig. 1A, *P* = 0.0089). The maximal effective i.t. morphine dose was 0.3 nmol. Over the time-course of treatment, morphine-induced itch responses mainly occurred in the first 30 min after injection. Intrathecal morphine-induced itch was significantly blocked under pretreatment with the MOR selective antagonist, CTOP, which suggests that morphine-induced itch response is mediated by MOR (Fig. 1B, *P* = 0.0153). Next, we examined whether morphine-induced itch is mediated by excitatory or inhibitory neurons in the spinal cord. To this end, we used *Vgat-Cre* or *Sst-Cre* mice crossed with *Oprm1*^*fl/fl*^ mice to conditionally knockout MOR on inhibitory or excitatory interneurons (Chamessian *et al*., 2018; Duan *et al*., 2018; Huang *et al*.) in the spinal cord, respectively. Strikingly, conditional deletion of *Oprm1* on inhibitory interneurons completely abolished morphine-induced itch (Fig. 1C. *P* = 0.0094). In sharp contrast, conditional deletion of *Oprm1* on excitatory interneurons did not affect morphine-induced itch behavior (Fig. 1D. *P* = 0.4476). These results indicate that i.t. morphine acts on MOR on inhibitory interneurons to evoke itch response. To confirm the specific involvement of MOR in this process, we tested scratching behavior induced by DAMGO, a MOR-selective agonist. Intrathecal DAMGO induced scratching in a bell-shaped dose response curve, and pruritus mainly occurred in the first 5 min (Fig. 1E). Like morphine, i.t. DAMGO-induced itch was significantly abolished in mice with conditional knockout of MOR on inhibitory interneurons (Fig. 1F, *P* = 0.0075).

Intrathecally injected morphine can act on MOR found on DRG afferent terminals as well as on interneurons in the spinal cord. To distinguish between the roles of peripherally and centrally expressed MOR in the context of intrathecal morphine-induced itch, we used the opioid receptor antagonist naloxone methiodide, which is unable to cross the blood-brain barrier (BBB) (Tejada *et al*., 2017; Xu *et al*., 2020). Intraperitoneal pretreatment with naloxone methiodide, which only blocks MOR activity in the peripheral sensory neurons, did not alter morphine-induced scratching (Fig. 2A, *P* = 0.1873). This result is consistent with a previous study in non-human primates, supporting the predominant role of spinal MOR in opioid-induced itch (Ko *et al*., 2004). In contrast, i.t. pretreatment with naloxone methiodide totally abolished morphine-induced itch, indicating that central spinal MOR signaling mediates morphine-induced itch (Fig. 2B, *P* = 0.0004). This finding was confirmed by the observation of very mild changes in i.t. morphine-induced itch in mice with conditional knockout of MOR on TRPV1^+^ sensory neurons and in mice pretreated with resininferatoxin (RTX, 1mg/kg), which ablates TRPV1^+^ sensory neurons (Fig. 2C, D, *P* = 0.2005, *P* = 0.4951, respectively). Furthermore, intradermal injection of morphine (100 nmol) evoked a very mild itch response (average of 14 scratches in 30 min), and this response was not affected in *Vgat-Cre; Oprm1*^*fl/fl*^ mice (Fig. 2E, *P* = 0.3281). These data suggest that i.t. morphine-induced itch is specifically mediated by MOR on inhibitory interneurons in the spinal cord.

### Intrathecal morphine induces antinociception through MOR on both peripheral sensory neurons and spinal cord excitatory interneurons

Spinal delivery of opioids is widely used in the clinical setting to provide analgesia. We likewise tested i.t. morphine-induced antinociception in MOR conditional knockout mice. Firstly, conditional knockout of MOR in TRPV1^+^, Vgat^+^, and SST^+^ neurons does not change baseline tail-flick latency (Supplementary Fig. 1A, *P* > 0.9999, *P* = 0.1324, *P* = 0.2199, respectively) or baseline hot-plate paw withdrawal latency (Supplementary Fig. 1B, *P* = 0.7716, *P* > 0.9999, *P* > 0.9999, respectively). In tail-flick testing, i.t. morphine significantly increased tail-flick latency, and this antinociceptive effect was significantly reduced in *Sst-Cre; Oprm1*^*fl/fl*^ mice (Fig. 4A, *P* = 0.0016), but not in *Vgat-Cre; Oprm1*^*fl/fl*^ (Fig. 4A, *P* = 0.4893) or *Trpv1-Cre; Oprm1*^*fl/fl*^ mice (Fig. 4A, *P* = 0.0801), although there was a weak decrease observed in *Trpv1-Cre; Oprm1*^*fl/fl*^ mice. In hot-plate testing, i.t. morphine-induced antinociception was significantly decreased in both *Sst-Cre; Oprm1*^*fl/fl*^ mice and *Trpv1-Cre; Oprm1*^*fl/fl*^ mice (Supplementary Fig. 1C, *P* < 0.0001 for both). Intrathecal morphine-induced antinociception was significantly enhanced in *Vgat-cre; Oprm1*^*fl/fl*^ mice (Supplementary Fig. 1C, *P* = 0.0050). Taken together, these results indicate that i.t. morphine-induced antinociception is mediated by MOR on spinal cord excitatory interneurons and peripheral sensory neurons.

**Figure 3.**
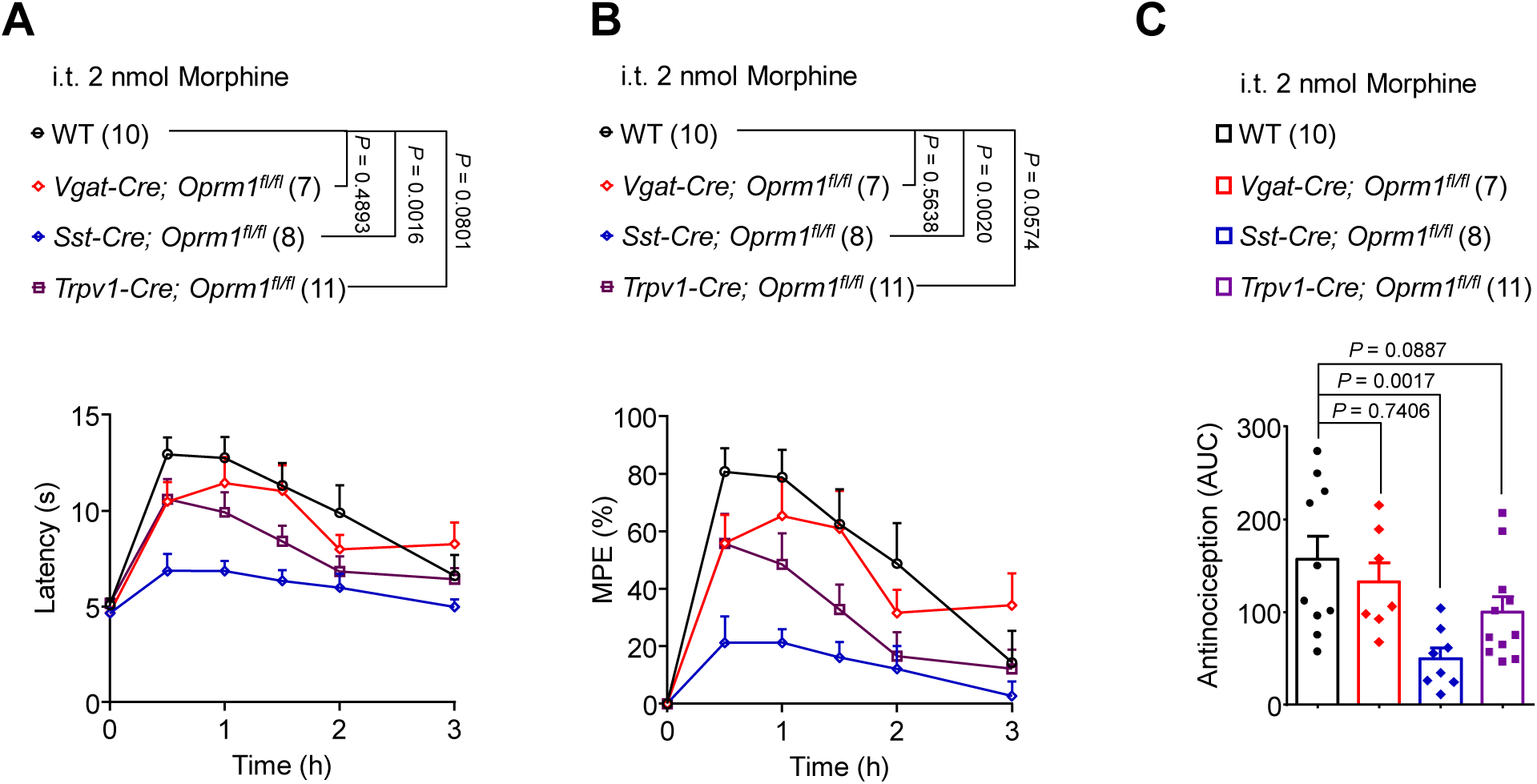
Intrathecal morphine-induced antinociception requires MOR expressed by spinal cord excitatory interneurons and peripheral sensory neurons. **(A)** Time course of the effect of intrathecal morphine (2 nmol) on tail-flick latency in WT, *Vgat-Cre; Oprm1*^*fl/fl*^, *Sst-Cre; Oprm1*^*fl/fl*^, and *Trpv1-Cre; Oprm1*^*fl/fl*^ mice. *P* = 0.4893, *P* = 0.0016, *P* = 0.0801, vs. WT mice, two-way ANOVA. **(B)** MPE of intrathecal morphine (2 nmol) induced antinociception in WT, *Vgat-Cre; Oprm1*^*fl/fl*^, *Sst-Cre; Oprm1*^*fl/fl*^, and *Trpv1-Cre; Oprm1*^*fl/fl*^ mice in the tail-flick test. *P* = 0.5638, *P* = 0.0017, *P* = 0.0574, vs. WT mice, two-way ANOVA. **(C)** AUC data converted from (B). *P* = 0.7406, *P* = 0.0017, *P* = 0.0887, vs. WT mice, one-way ANOVA. Data are mean ± SEM. Sample sizes are indicated in the figures.

**Figure 4.**
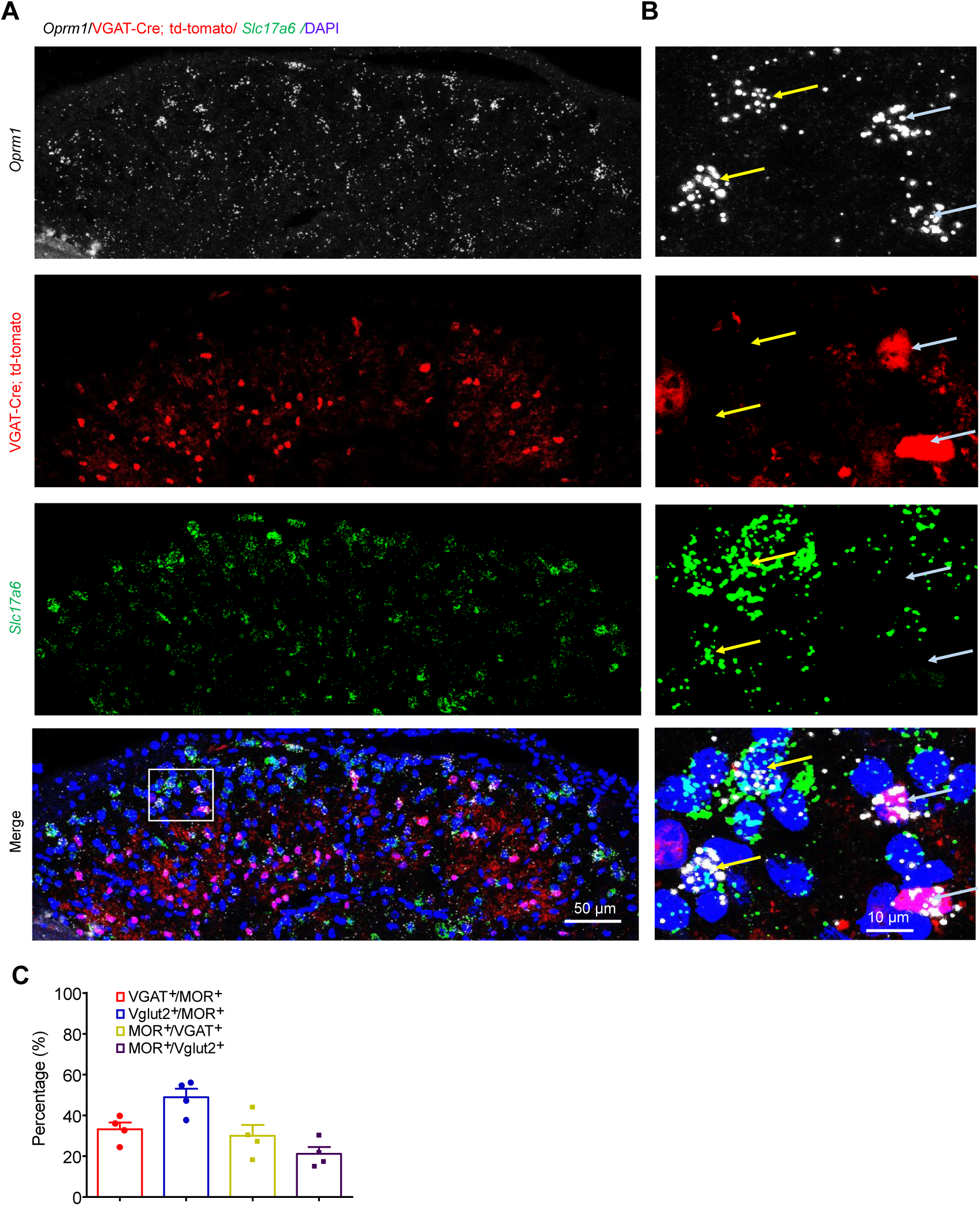
MOR is expressed by both inhibitory and excitatory interneurons in SDH. **(A, B)** *In situ* hybridization (ISH, RNAscope images of MOR mRNA (*Oprm1*, white) and Vglut2 mRNA (*Slc17a6*, green) in mouse SDH of *Vgat-Cre; tdTomato* mice. Yellow stars and white stars indicate *Oprm1* double-labeled with Vglut2 or Vgat, respectively. Scale bar, 50 μm (A) 10 μm (B). **(C)** The percentage of co-expression between MOR and Vgat or Vglut2 neurons in SDH. 8 spinal cord sections from 4 mice were analyzed. Data are mean ± SEM.

### MOR agonists suppress the activities of MOR^+^ inhibitory interneurons in spinal cord slices

We next examined MOR expression in the spinal dorsal horn (SDH) by RNAscope assay. *Oprm1* mRNA expression on excitatory and inhibitory neurons was tested by staining for *Oprm1* with Vglut2 mRNA (*Slc17a6*) in *Vgat*-Cre; Ai9 tdTomato mice. *Oprm1* expression was observed on both Vgat^+^ and Vglut2^+^ interneurons in lamina II of the SDH (Fig. 4A-C). This finding is in line with published single cell sequencing data which recorded broad MOR expression in both inhibitory and excitatory interneurons of the SDH (Haring *et al*., 2018). However, the expression level of MOR in inhibitory interneurons is higher than in excitatory interneurons (Supplementary Fig. 5A) (Haring *et al*., 2018)

Next, we performed *ex vivo* electrophysiology recordings of spinal cord slices to evaluate the effects of MOR agonists on the activities of inhibitory interneurons, taking note that opioids activate potassium channels through G protein-coupled receptors to generate outward currents in MOR expressing neurons (North and Williams, 1985). Our results showed that DAMGO (0.5 μM) evoked outward currents in 9 out of 19 SDH lamina II inhibitory interneurons with an amplitude of 19.5 pA in Vgat^+^ interneurons within spinal cord slices from *Vgat-Cre*; Ai9 tdTomato reporter mice (Fig. 5A). We also recorded the effects of morphine on current injection-evoked action potentials in SDH lamina II Vgat^+^ interneurons from *Vgat-Cre*; Ai9 reporter mice. Perfusion of morphine (10 μM) significantly inhibited evoked action potentials in SDH lamina II inhibitory interneurons (Fig. 5B and C, *P* = 0.0109).

**Figure 5.**
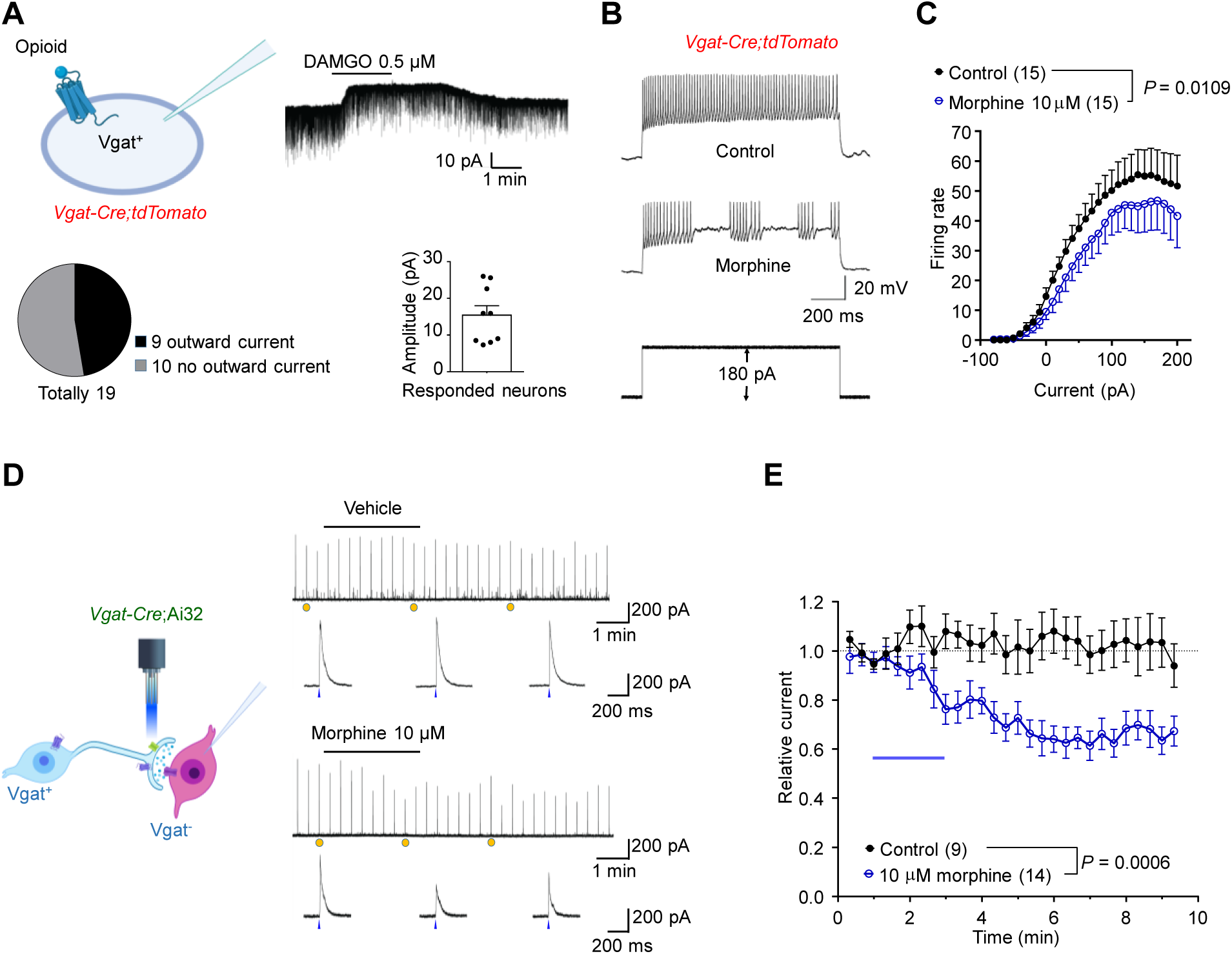
DAMGO and morphine inhibit the activities of MOR^+^ inhibitory interneurons in SDH of spinal cord slices. **(A)** DAMGO (0.5 µM) induced outward currents recorded from Vgat^+^ interneurons in the SDH from *Vgat-Cre; tdTomato* mice. 9 of 19 recorded neurons showed outward currents with an average current of 19.50 pA. **(B)** Typical trace of action potentials from Vgat^+^ interneurons in the spinal cord dorsal horn from *Vgat-Cre; tdTomato* mice. **(C)** Quantification of the effects of morphine (10 µM) on action potentials, *P* = 0.0109, two-way ANOVA. **(D)** Traces of light-evoked IPSCs in the spinal cord from *Vgat-Cre; Ai32* mice treated with vehicle (upper) or morphine (bottom). **(E)** Quantification of morphine (10 µM) effects on evoked IPSC, *P* = 0.0006, two-way ANOVA. Data are mean ± SEM. Sample sizes are indicated in the figures.

To further characterize the functional inhibition of inhibitory interneurons in neurotransmission, we expressed channelrhodopsin ChR2 in spinal Vgat^+^ interneurons by crossing *Vgat*-Cre mice with Ai32 mice. Evoked inhibitory postsynaptic currents (IPSCs) in lamina II Vgat^-^ interneurons were recorded while spinal Vgat^+^ interneurons were stimulated by application of 473 nm wavelength blue light. Morphine (10 μM) significantly inhibited evoked IPSCs, indicating a functional inhibition of inhibitory interneurons (Fig. 5D and E, *P* < 0.0001). These results demonstrate that µ-opioids can directly inhibit the activities of MOR^+^ inhibitory interneurons, suggesting that opioid-induced itch is a result of inhibition of MOR^+^ inhibitory interneurons.

### Intrathecal morphine induced itch is suppressed by NPY and abolished by ablation of GRPR^+^ interneurons

Two populations of SDH inhibitory interneurons, namely NPY^+^ and dynorphin (DYN)^+^ interneurons, have been implicated in the gate control of itch transmission (Ross *et al*., 2010; Kardon *et al*., 2014; Bourane *et al*., 2015; Acton *et al*., 2019; Pan *et al*., 2019). We first examined MOR expression in these two populations of inhibitory interneurons by triple ISH for *Oprm1, Npy* and *Pdyn* mRNA expression in the SDH. We found that MOR in inhibitory interneurons is mainly expressed in NPY^+^ interneurons (Supplementary Fig. 2A and B). The level of co-expression between MOR and NPY is similar to that of MOR and Vgat, which indicates MOR in inhibitory interneurons is mainly co-expressed with NPY. This is consistent with data from a single-cell sequencing database (Supplementary Fig. 5) (Haring *et al*., 2018). Notably, i.t. co-administration of NPY (10 µg) with morphine (0.3 nmol) significantly inhibited morphine-induced itch (Fig. 6A, *P* < 0.0001). Spinal NPY-Y1 receptor is involved in itch transmission (Acton *et al*., 2019). We observed that Y1R is highly co-expressed with GRP (70%) (Supplementary Fig. 3). The expression patterns of MOR, NPY, and Y1 together suggest that MOR activation in NPY^+^ inhibitory interneurons may disinhibit GRP^+^ interneurons to produce itch. To test this hypothesis, we intrathecally injected bombesin-saporin (400 ng) to ablate GRPR^+^ interneurons in SDH, as previously demonstrated (Liu *et al*., 2011; Pan *et al*., 2019). In bombesin-saporin pretreated mice, i.t. GRP (0.1 nmol)-induced pruritus was abolished, indicating successful ablation of GRPR^+^ interneurons (Fig. 6B, *P* = 0.0049). Notably, i.t. morphine-induced itch was totally abolished by ablation of GRPR^+^ neurons, suggesting an essential role of GRPR^+^ interneurons in morphine-induced itch (Fig. 6C, *P* = 0.0072). In contrast, intrathecal GRP-induced itch was not affected in *Vgat-Cre; Oprm1*^*fl/fl*^ mice (Fig. 6D, *P* = 0.7797). RNAscope data revealed that MOR has low levels of co-expression with GRP^+^ interneurons (29.4%) and GRPR^+^ interneurons (4.7%) (Supplementary Fig. 4), which is in agreement with single-cell sequencing data which shows very low expression levels of MOR in GRP^+^ and GRPR^+^ neurons (Supplementary Fig. 5) (Haring *et al*., 2018). Taken together these data indicate that the action of GRP/GRPR lies downstream of MOR^+^ inhibitory interneurons in the production of i.t. morphine-induced itch.

**Figure 6.**
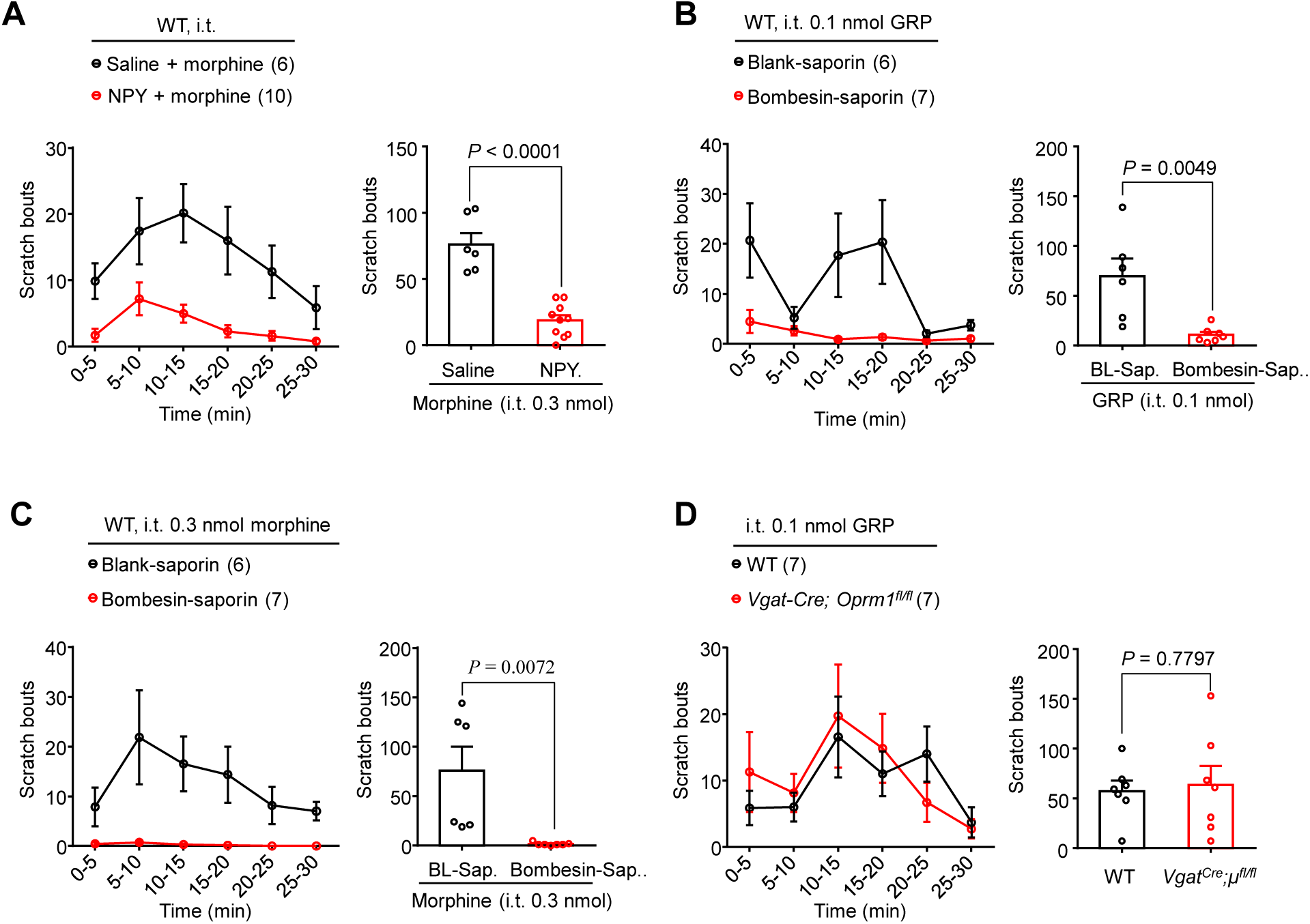
Morphine-induced itch is suppressed by intrathecal NPY and ablation of GRPR^+^ neurons. **(A)** Intrathecal morphine (0.3 nmol) induced itch is inhibited by co-injection of NPY (10 µg, i.t.) in WT mice. *P* < 0.0001, two-tailed Student’s t-test. **(B)** Intrathecal GRP (0.1 nmol) induced itch is abolished by ablation of spinal GRPR^+^ neurons via intrathecal injection of Bombesin-saporin (400 ng). *P* = 0.0049, two-tailed Student’s t-test. **(C)** Intrathecal morphine (0.3 nmol) induced itch is abolished in Bombesin-saporin treatment. *P* = 0.0072, two-tailed Student’s t-test. **(D)** Intrathecal GRP (0.1 nmol) induced itch is not affected in *Vgat-Cre; Oprm1*^*fl/fl*^ mice. *P* = 0.7797, two-tailed Student’s t-test. Data are mean ± SEM.

### MOR in inhibitory interneurons contributes to dermatitis-associated persistent itch but not to acute chemical itch

We further investigated the contribution of MOR to itch in different animal models of pruritus. We first used a DNFB-induced allergic contact dermatitis model (Liu *et al*., 2016) to determine the roles of MOR in chronic itch. DNFB-induced itch was significantly reduced in *Vgat-Cre; Oprm1*^*fl/fl*^ mice (Fig. 7A and B, *P* = 0.0224). Moreover, we tested acute itch induced by chloroquine (CQ) and histamine. Acute itch induced by intradermal injection of CQ and histamine did not significantly change in *Vgat-Cre; Oprm1*^*fl/fl*^ mice (Fig. 7C and D, *P* = 0.6328, *P* = 0.5120, respectively). These results indicate that MOR on inhibitory interneurons specifically contributes to dermatitis-associated chronic itch but does not have an active role in acute chemical itch.

**Figure 7.**
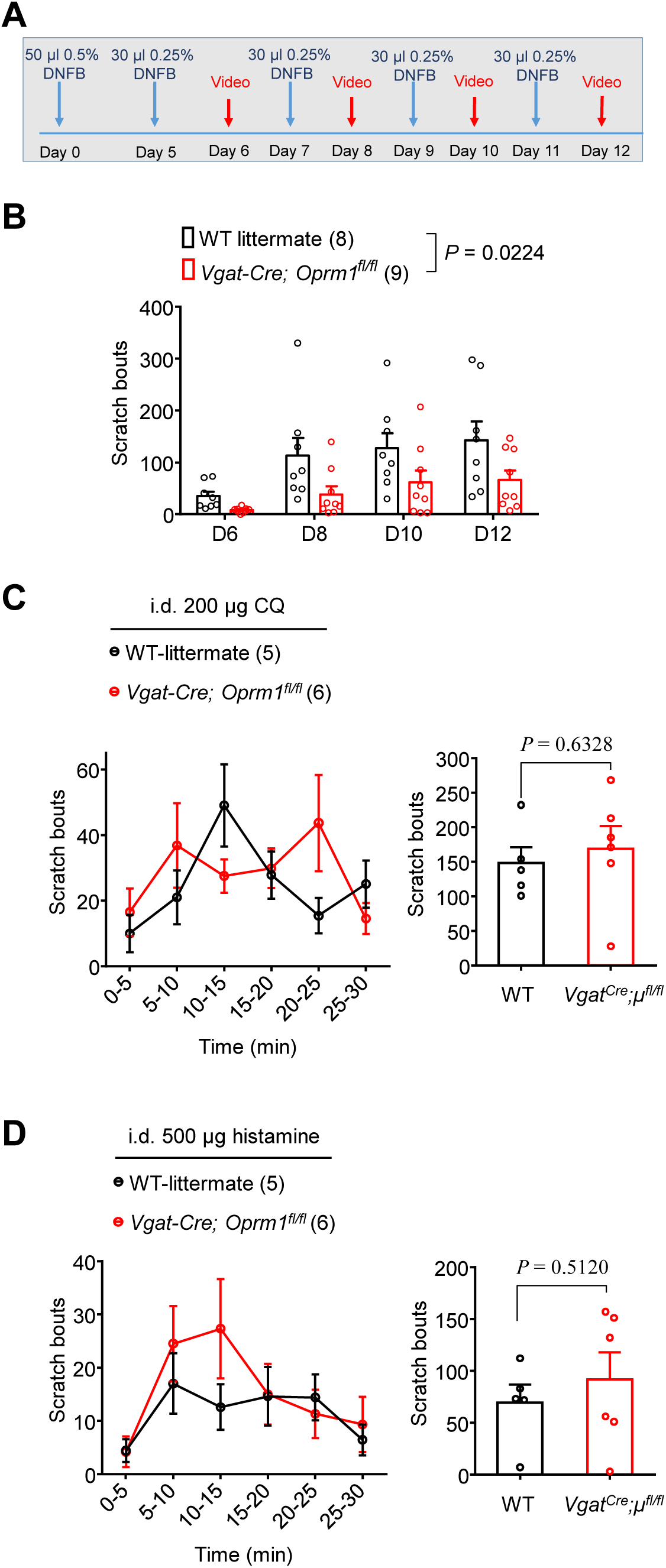
MOR on inhibitory neurons regulates chronic itch but not acute itch. **(A)** Protocol of experimental design for the DNFB mouse model. **(B)** Spontaneous itch in the DNFB mouse model is significantly decreased in *Vgat-Cre; Oprm1*^*fl/fl*^ mice. *P* = 0.0224, two-way ANOVA. **(C)** Time course (left) and total scratch bouts (right) within 30 min after intradermal injection of 200 µg CQ in WT and *Vgat-Cre; Oprm1*^*fl/fl*^ mice. *P* = 0.6328, two-tailed Student’s t-test. **(D)** Time course (left) and total scratch bouts (right) within 30 min after intradermal injection of 500 µg histamine in WT and *Vgat-Cre; Oprm1*^*fl/fl*^ mice. *P* = 0.6328, two-tailed Student’s t-test. Data are mean ± SEM. Sample sizes are indicated in the figures.

### Centrally but not peripherally restricted MOR antagonist inhibits dermatitis and lymphoma-induced chronic itch

We used the peripherally-restricted opioid receptor antagonist naloxone methiodide to assess the role of peripherally expressed MOR in dermatitis-induced persistent itch. Intraperitoneal injection of naloxone but not naloxone methiodide significantly reduced spontaneous itch in the DNFB model suggesting that MOR expressed in the CNS more critically involved in chronic itch (Fig 8A, *P* = 0.0044 and *P* = 0.6383, respectively). Furthermore, i.t. naloxone significantly reduced DNFB-induced itch (Fig. 8B, *P* = 0.0007).

**Figure 8.**
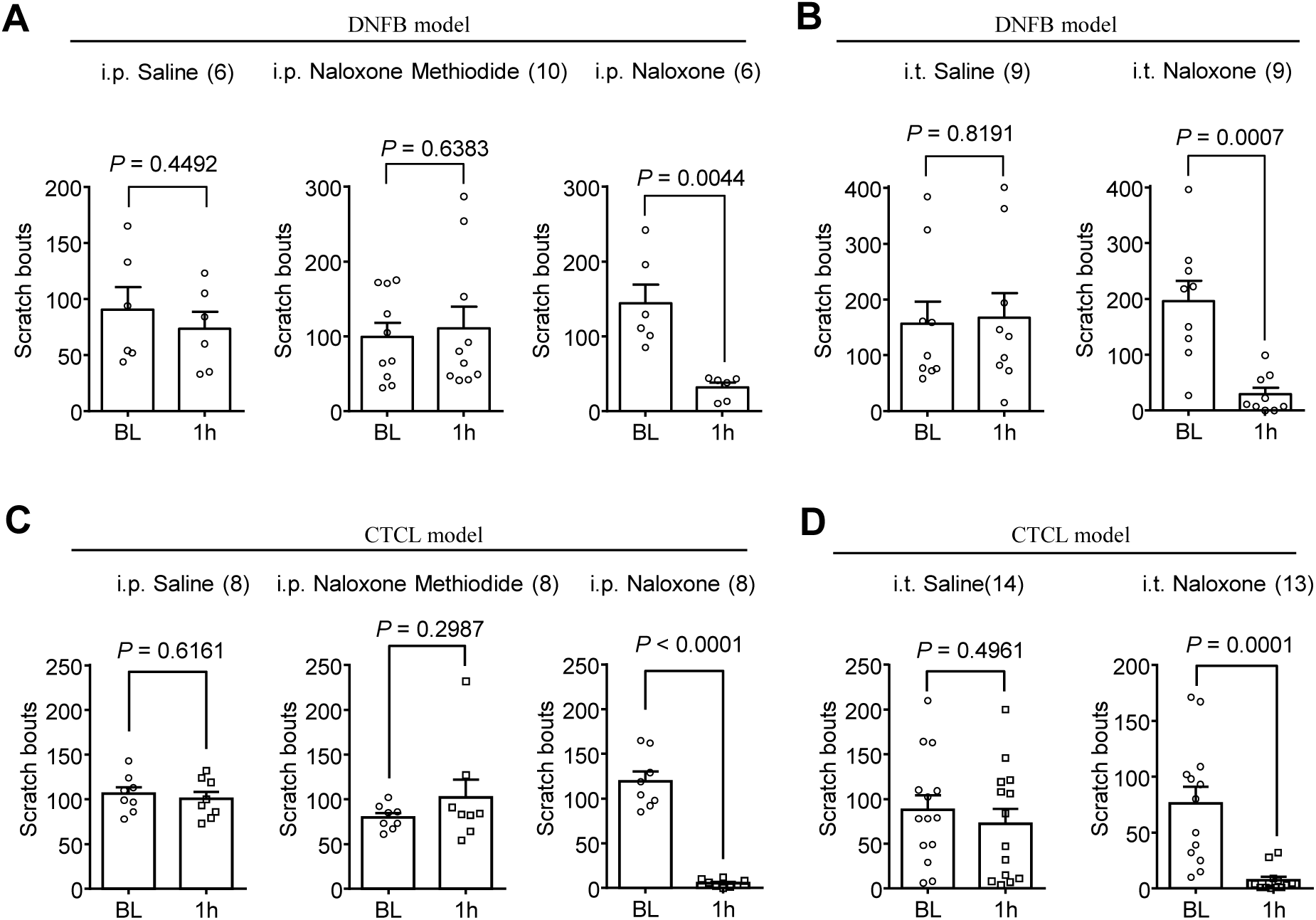
Central MOR contributes to chronic itch in the DNFB model and CTCL model. **(A)** Effects of intraperitoneal pretreatment with saline, naloxone methiodide, or naloxone on DNFB-induced itch. *P* = 0.4492, *P* = 0.6383, and *P* = 0.0044, respectively, two-tailed Student’s t-test. **(B)** The effects of i.t. pretreatment with saline or naloxone on DNFB-induced spontaneous. *P* = 0.8191 and *P* = 0.0007, respectively, two-tailed Student’s t-test. **(C)** The effects of intraperitoneal pretreatment with saline, naloxone methiodide, or naloxone, on spontaneous itch in the CTCL model. *P* = 0.6161, *P* = 0.2987, and *P* < 0.0001, respectively, two-tailed Student’s t-test. **(D)** The effects of intrathecal pretreatment with saline, or naloxone on CTCL-induced chronic itch. *P* = 0.4961 and *P* = 0.0001, respectively, two-tailed Student’s t-test. Data are mean ± SEM. Sample sizes are indicated in the figures.

Finally, we tested opioid antagonists in a mouse cutaneous T-cell lymphoma (CTCL) model, which was recently developed in our lab by intradermal inoculation of human Myla cells (Han *et al*., 2018). In the CTCL model, mice develop lymphoma and chronic itch (Han *et al*., 2018). Similar to the results found in the DNFB model, only systemic treatment with naloxone but not peripherally restricted naloxone methiodide significantly inhibited spontaneous itch in CTCL mice (Fig. 8C, *P* < 0.0001 and *P* = 0.2987, respectively). Intrathecally injected naloxone also significantly abolished spontaneous itch in the CTCL model (Fig. 8D, *P* = 0.0001). These results indicate that centrally but not peripherally expressed MOR plays a major role in dermatitis and lymphoma-induced chronic itch.

## Discussion

Itch is a common side effect of opioids when used as a clinical treatment for pain. In this study, we demonstrated that intrathecal injection of the µ-opioids morphine and DAMGO elicited itch responses that are mainly mediated by MOR on spinal GABAergic inhibitory interneurons. µ-opioids can inhibit the activities of Vgat^+^ interneurons in the spinal cord and disinhibit the itch signaling pathway, resulting in the generation of itch behavior. Additionally, chronic itch in the DNFB-induced allergic contact dermatitis mouse model was decreased after *Oprm1*-*Vgat* deletion. Finally, naloxone, but not peripherally-restricted naloxone methiodide, inhibited itch in both the DNFB and CTCL models, indicating the contribution of central MORs to chronic itch. Our findings demonstrate that i.t. opioid acts upon MORs on spinal inhibitory interneurons to regulate pruritus via disinhibition. Our data also suggest that chronic itch could be effectively treated with CNS-targeted naloxone.

Recent progress has advanced our understanding regarding the mechanisms of itch (LaMotte *et al*., 2014; Ji, 2018; Cevikbas and Lerner, 2019). The itch sensation is initiated by the activation of distinct populations of primary pruriceptors expressing MAS-related GPR receptors (Mrgpr), histamine receptors, 5-HT receptors, and natriuretic peptide B (Nppb) by a variety of pruritogens (Liu *et al*., 2009; Han *et al*., 2013; Mishra and Hoon, 2013; Pan *et al*., 2019). The pruriceptors transmit itch signals to the spinal cord by release of neuropeptides such as Nppb in the SDH, but also see (Sun and Chen, 2007). The spinal interneurons expressing natriuretic peptide receptor 1 (NPR1) or GRP receive and process chemical itch signals (Sun and Chen, 2007; Sun *et* al., 2009; Mishra and Hoon, 2013; LaMotte *et al*., 2014). Like mechanical allodynia, mechanical itch could be mediated by a subtype of Aβ low-threshold mechanoreceptors expressing Toll-like receptor 5 (TLR5), transmitting itch signals to a subpopulation of excitatory interneurons expressing Urocortin 3 or NPY1R (Pan *et al*., 2019; Wang *et al*., 2019). Furthermore, two populations of inhibitory interneurons expressing Bhlhb5^+^ and NPY^+^ in the SDH are involved in modulating chemical and mechanical itch, respectively (Ross *et al*., 2010; Bourane *et al*., 2015; Acton *et al*., 2019; Pan *et al*., 2019). Despite this progress, the specific roles of MOR in itch signaling pathways remained unclear.

Intrathecal morphine induces paradoxical itch and analgesia, and different populations of projection neurons (e.g., trigeminothalamic tract neurons) are involved in i.t. morphine-induced itch and analgesia (Moser and Giesler, 2013). Pain is known to suppress itch via spinal thalamic tract neurons (Davidson *et al*., 2009). Our findings showed that i.t. morphine-evoked itch and antinociception is mediated by different populations of interneurons, specifically excitatory interneurons for antinociception and inhibitory neurons for pruritus. Loss of MOR in excitatory neurons (SST^+^) resulted in a substantial reduction (80%) in morphine-induced antinociception. Of interest, loss of MOR in TRPV1^+^ neurons only showed a partial reduction of i.t. morphine antinociception. Mechanistically, morphine binds to MOR on excitatory interneurons suppressing activity by inducing outward currents through inwardly rectifying potassium channels (North and Williams, 1985; Andrade *et al*., 2010). Morphine also inhibits neurotransmitter release in SDH neurons by inhibiting calcium channels on primary sensory neurons (Wang *et al*., 2020). Although GRPR^+^ neurons have been implicated as projection neurons, recent studies indicate these neurons are excitatory interneurons connected to NK1^+^ projection neurons (Acton *et al*., 2019; Bardoni *et* al., 2019). Consistently, itch is abolished by ablation of NK1^+^ projection neurons (Carstens *et al*., 2010). Liu et al found that morphine can activate GRPR^+^ to elicit itch through MOR1D by forming a heterodimer with GRPR in the SDH (Liu *et al*., 2011). In agreement with this study, we found that ablation of GRPR^+^ neurons totally blocked morphine-induced itch. However, our findings also suggested a distinct mechanism by which morphine acts on inhibitory neurons to elicit itch through disinhibition.

The most striking observation of this study is that conditional deletion of *Oprm1* in Vgat^+^ inhibitory neurons completely abolished pruritus induced by i.t. morphine or DAMGO. Our ISH data, in agreement with single-cell sequencing (Haring *et al*., 2018), demonstrated high expression of *Oprm1* in inhibitory neurons but almost no co-expression of *Oprm1* and *Grpr* in the SDH. Furthermore, our electrophysiological data confirmed that MOR agonists directly act on Vgat^+^ inhibitory neurons to induce outward currents and suppress action potentials on inhibitory neurons (Fig 5A-C) and produce sustained inhibition of IPSCs on postsynaptic Vgat^-^ excitatory neurons resulting in disinhibition (Fig. 5D-E). NPY was reported to modulate mechanical itch and histamine-induce itch (Gao *et al*., 2018; Acton *et al*., 2019; Pan *et al*., 2019), but it remained unclear if NPY contributes to morphine-induced itch. Our ISH and behavioral data demonstrated that 1) *Oprm1* is co-expressed with both Vgat and NPY, 2) i.t. NPY effectively inhibited morphine-induced itch, and 3) NPY1R is highly co-expressed with GRP in SDH neurons. Thus, we postulate that morphine activates MOR in NPY^+^ inhibitory interneurons to disinhibit GRP^+^ excitatory interneurons leading to the induction of itch. Previous studies showed that kappa opioid receptor also contributed to morphine-induced itch in primates (Ko and Husbands, 2009; Lee and Ko, 2015). Our ISH data revealed a moderate co-expression of *Pdyn* with *Oprm1* in mouse SDH. Thus, we should not rule out the involvement of *Pdyn*^+^ inhibitory neurons in morphine-induced itch. Further identification of opioid receptor-containing inhibitory interneuron subtypes in opioid-induced itch remains an important area for future research.

Patients with chronic itch commonly experience high sensitivity to pruritogens, mechanically evoked itch sensations, and spontaneous itch (Ikoma *et al*., 2006; LaMotte *et al*., 2014). Opioid receptor antagonists including naloxone, naltrexone, and nalbuphine have been demonstrated to be the effective treatments for chronic itch under certain pathological conditions (Brune *et al*., 2004; Reszke and Szepietowski, 2018; Serrano *et al*., 2018; Kremer, 2019). MOR antagonists has been shown to be effective treatments for dermatitis-associated itch (Monroe, 1989; Metze *et al*., 1999; Farmer and Marathe, 2017; Pavlis and Yosipovitch, 2018; Ekelem *et al*., 2019). Naltrexone is an effective treatment for reducing uremic pruritus in patients with chronic kidney disease (Peer *et al*., 1996; Legroux-Crespel *et al*., 2004). Furthermore, case reports have shown that naloxone and naltrexone are effective therapies for anti-PD1 immunotherapy-induced pruritus (Kwatra *et al*., 2018; Singh *et al*., 2019). Although MOR antagonists have shown some benefits for treating pruritus in these pathological conditions, the specific cellular targets of MOR antagonists for treating chronic itch remained unclear. Our present study revealed that persistent itch in DNFB-induced allergic contact dermatitis is significantly impaired under the conditional deletion of MOR in GABAergic neurons. Furthermore, naloxone but not peripherally-restricted naloxone methiodide effectively alleviated chronic itch in DNFB-induced allergic contact dermatitis and T cell-induced lymphoma mouse models. Further clinical studies will be needed to test the effects of intrathecal or CNS-penetrating MOR antagonists in patients suffering from various chronic itch conditions.

## Abbreviations

ACD: allergic contact dermatitis;
AUC: area under the curve;
CNS: central nervous system;
CKO: conditional knockout mice;
CQ: chloroquine;
CTCL: cutaneous T-cell lymphoma;
DNFB: 1-Fluoro-2,4-dinitrobenzene;
DYN: dynorphin;
GRP: gastrin releasing peptide;
GRPR: gastrin releasing peptide receptor;
INs: interneurons;
IPSC: inhibitory postsynaptic current;
ISH: In situ hybridization;
MOR: mu opioid receptor;
MPE: maximum possible effect;
NPY: neuropeptide Y;
PNS: peripheral nervous system;
RTX: resiniferatoxin;
SDH: spinal cord dorsal horn;
SOM: somatostatin;
TRPV1: transient receptor potential ion channel subtype V1

## Supplementary material

**Figure S1.**
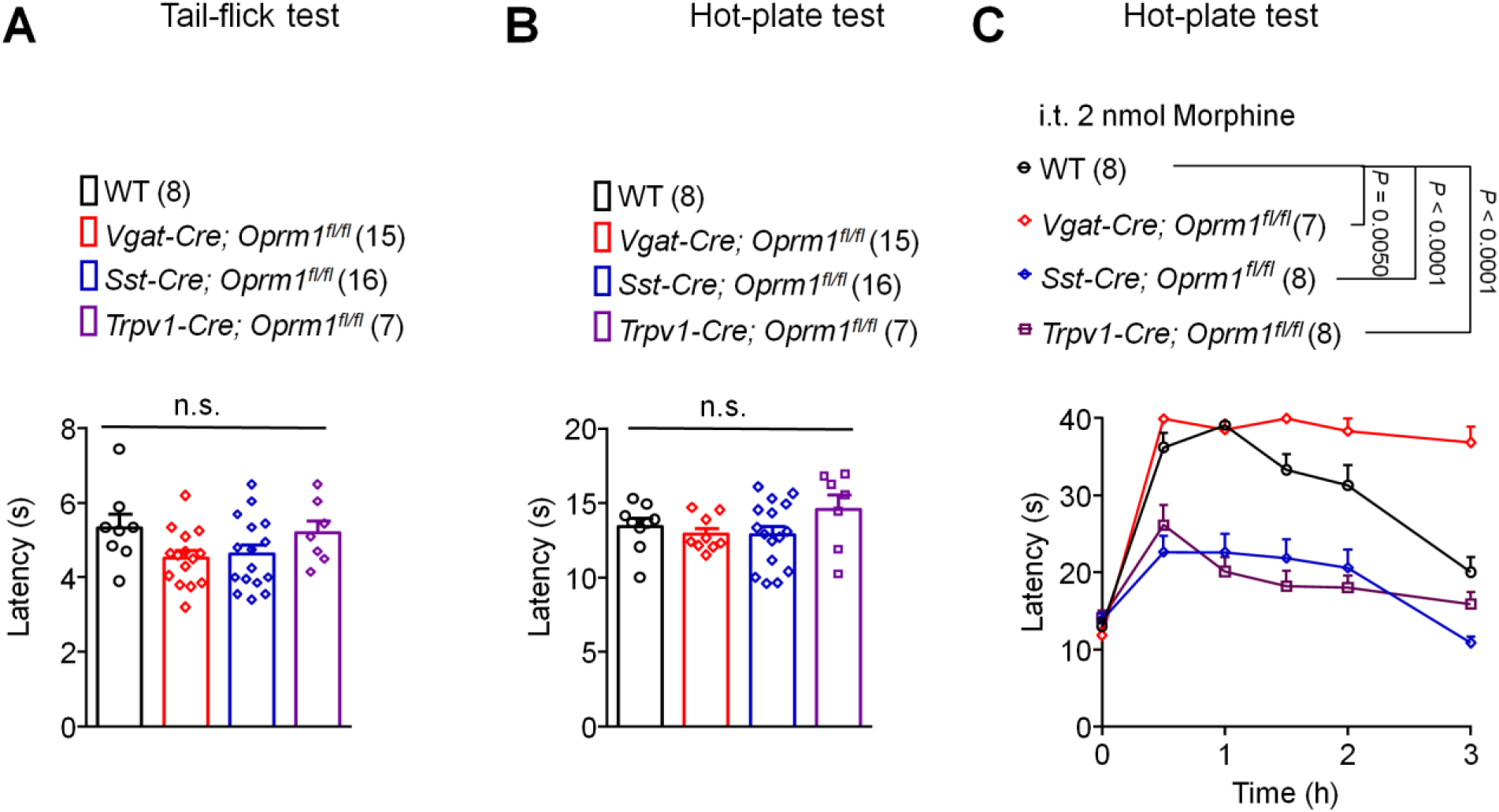
Tail-flick and hot-plate tests in wild-type and transgenic mice. **(A-B)** Tail-flick (A) and hot-plate (B) tests in WT, *Vgat-Cre; Oprm1*^*fl/fl*^, *Sst-Cre; Oprm1*^*fl/fl*^, *Trpv1-Cre; Oprm1*^*fl/fl*^ mice. n.s., no significance, one-way ANOVA. **(C)** Hot plate testing for intrathecal 2 nmol morphine induced antinociception in WT, *Vgat-Cre; Oprm1*^*fl/fl*^, *Sst-Cre; Oprm1*^*fl/fl*^, *Trpv1-Cre; Oprm1*^*fl/fl*^ mice. *P* = 0.0050, *P* < 0.0001, *P* < 0.0001, vs WT mice, two-way ANOVA. Data are Mean ± SEM. Sample sizes are indicated in the figures.

**Figure S2.**
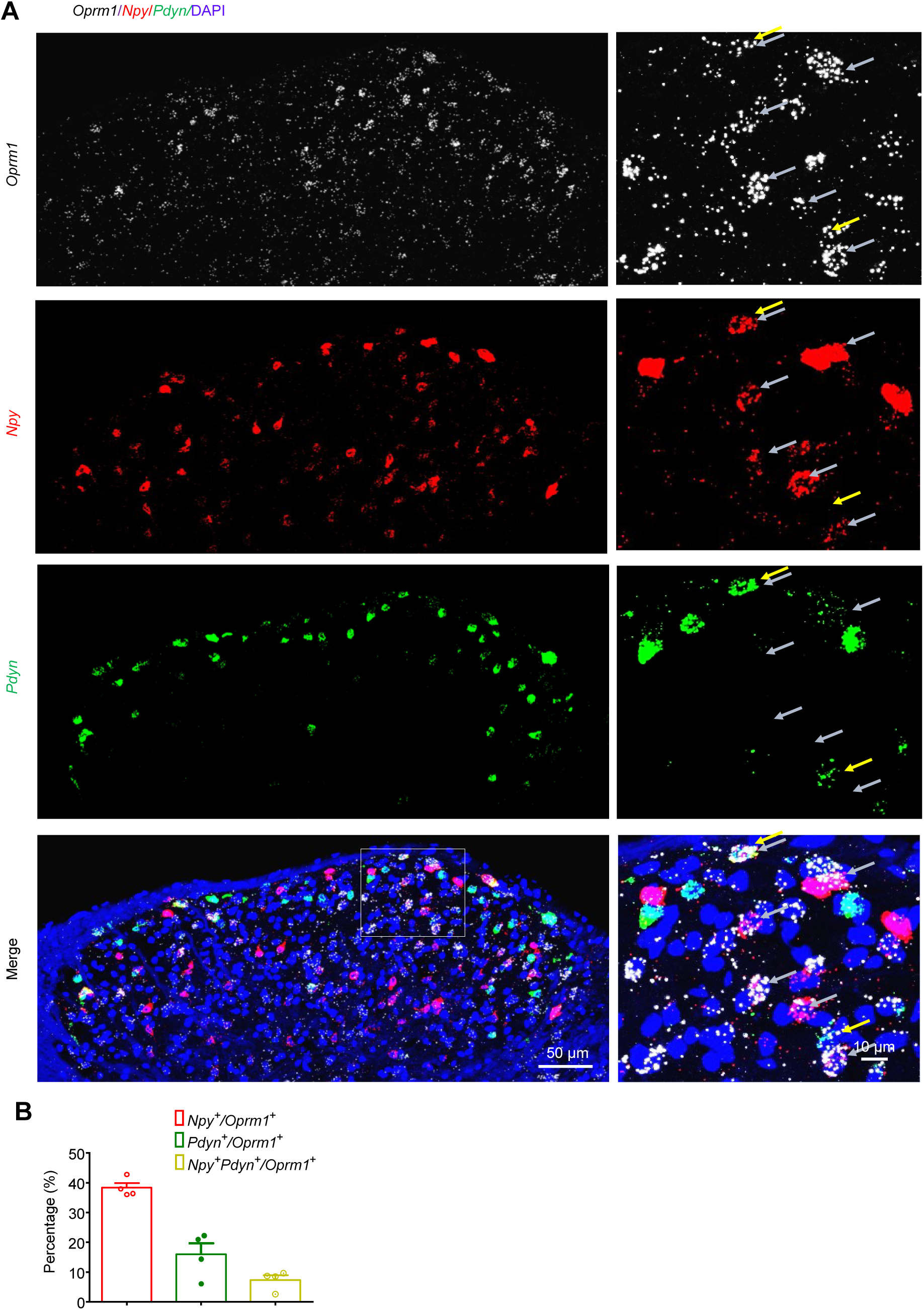
MOR expression in NPY^+^ inhibitory interneurons in the spinal cord dorsal horn (SDH). **(A)** In situ hybridization (RNAscope) images showing mRNA expression for *Oprm1* (white), *Npy* (red) and *Pdyn* (encoding Pro-Dynorphin, green) in WT mouse SDH. Yellow and gray stars indicate *Oprm1* co-localization with *Pdyn* or *Npy*, respectively. Scale bar, 50 μm (left) 10 μm (right). **(B)** The percentage of co-expression of *Oprm1* with *Npy* and *Pydn* in SDH neurons. Eight spinal cord sections from 4 mice were analyzed.

**Figure S3.**
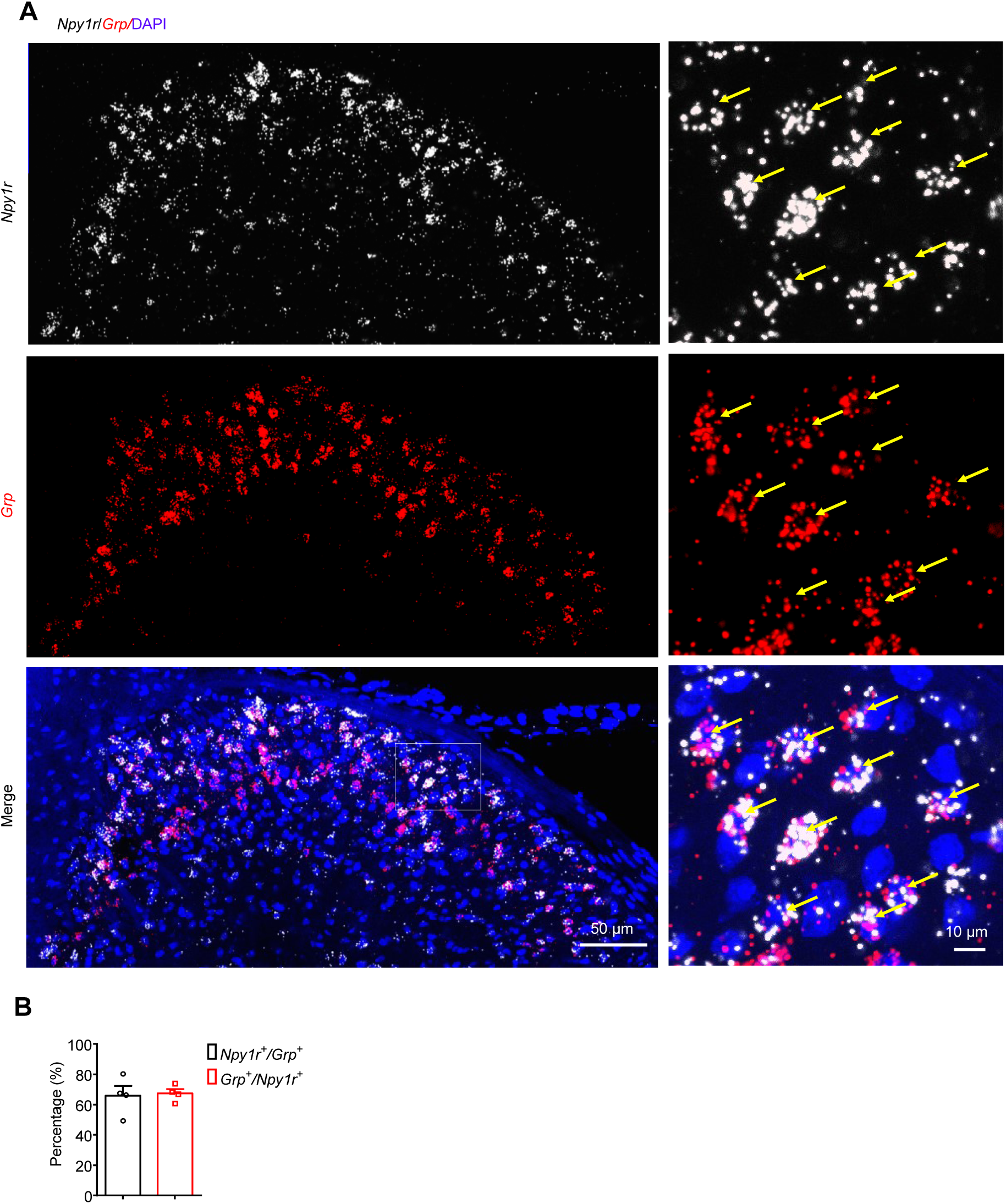
Co-expression of GRP and NPY1R in the SDH. **(A)** In situ hybridization (RNAscope) images showing the co-expression of *Npy1r* (white) and *Grp* (red) in WT mouse SDH. Yellow stars show *Npy1r* double labelling with *Grp*. Scale bar, 50 μm (left) 10 μm (right). **(B)** The percentage of co-expression of *Npy1r* with *Grp* in SDH neurons. Eight spinal cord sections from 4 mice were analyzed. Data are Mean ± SEM.

**Figure S4.**
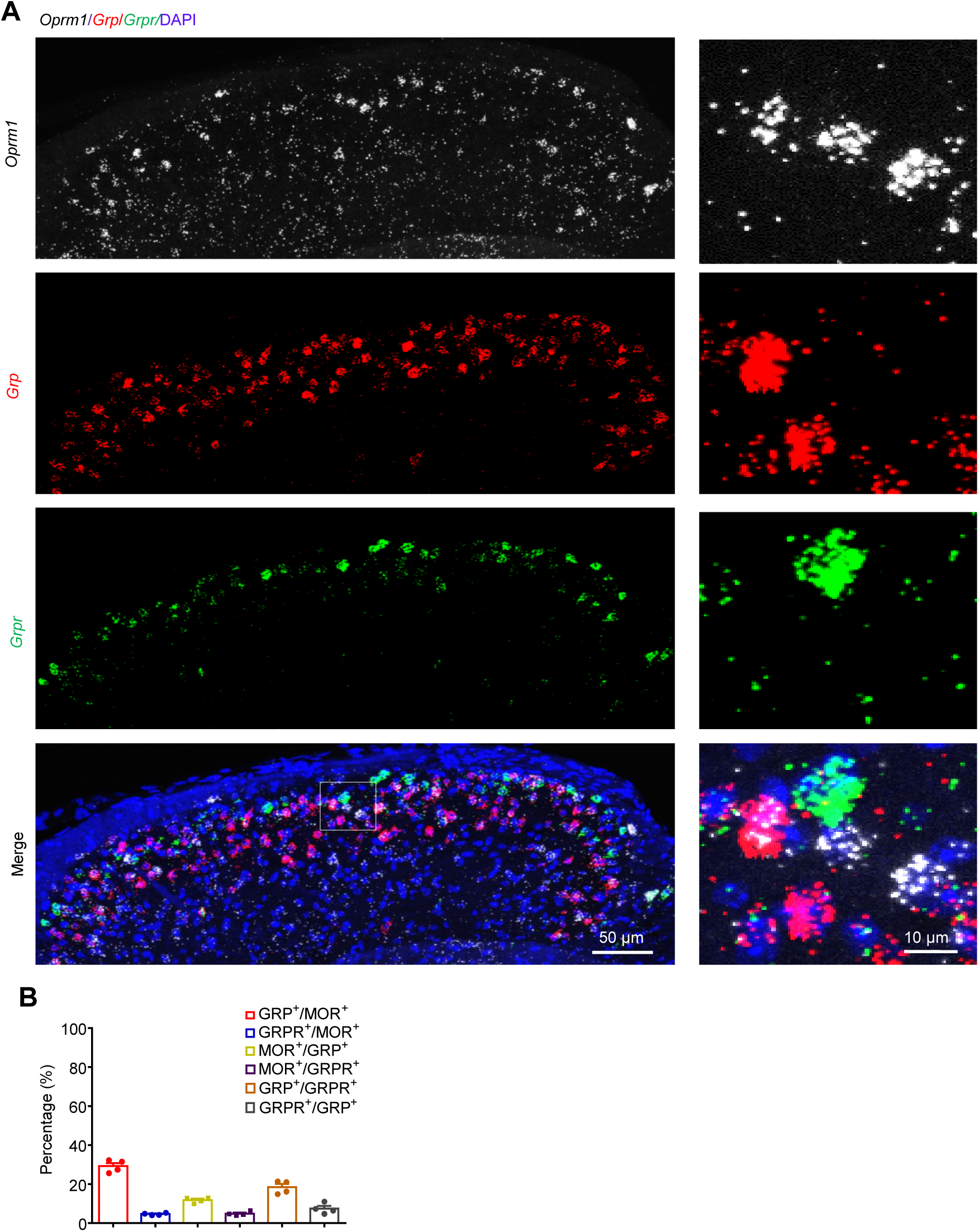
Limited co-expression of MOR with GRPR in the SDH. **(A)** In situ hybridization (RNAscope) images showing mRNA expression of *Oprm1* (white), *Grp* (red), and *Grpr* (green) in WT mouse SDH. Scale bar, 50 μm (left) 10 μm (right). **(B)** The percentage of co-expression of MOR^+^, GRP^+^, GRPR^+^ neurons in mouse SDH. Eight spinal cord sections from 4 mice were analyzed. Data are Mean ± SEM.

**Figure S5.**
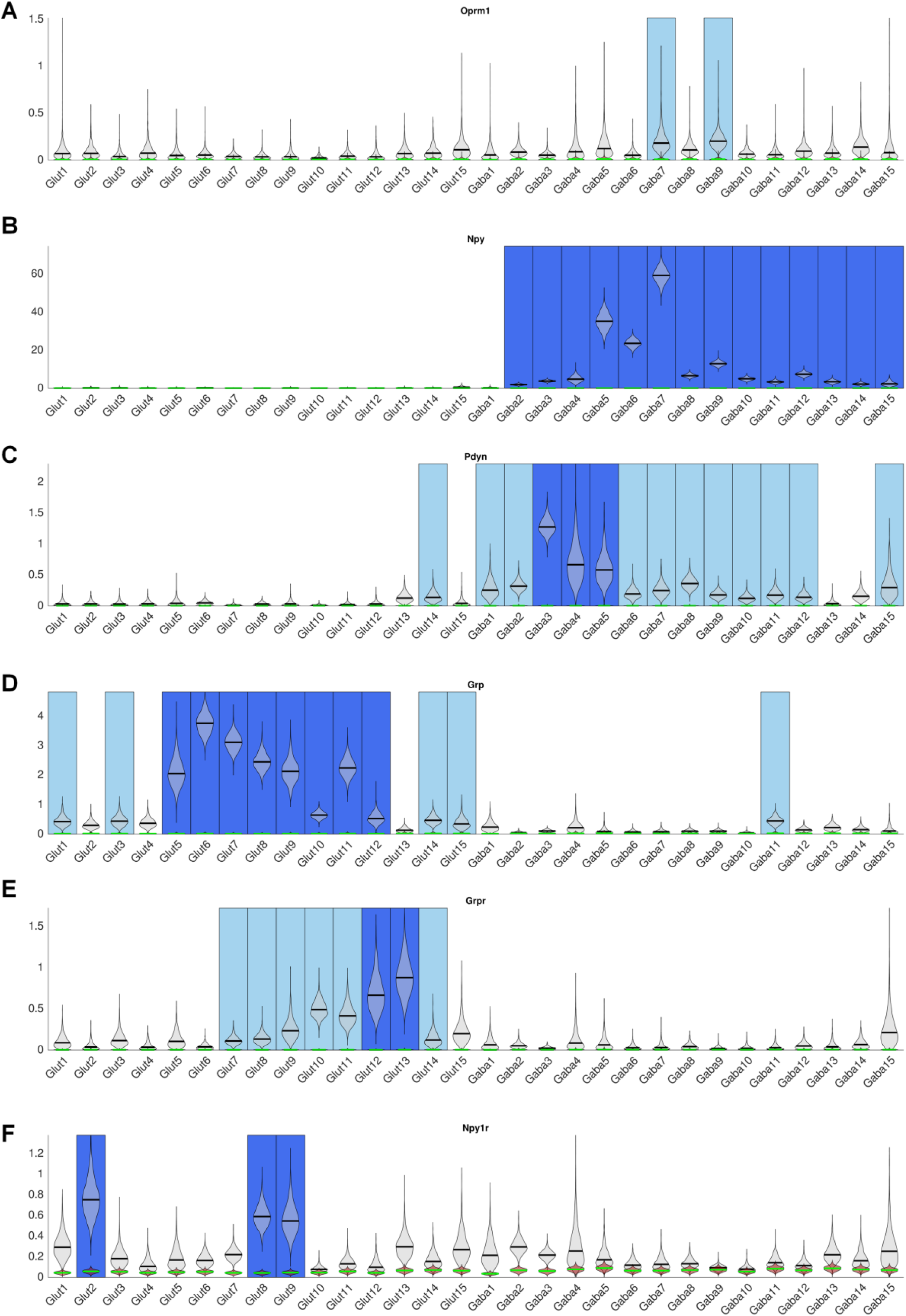
Single-cell RNA sequencing database showing gene expression in mouse SDH. Expression patterns of *Oprm1* (A), *Npy* (B), *Pdyn* (C), *Grp* (D), *Grpr* (E), and *Npy1r* (F) in the mouse spinal cord from the single cell RNA sequencing data (Cited from Haring *et al*., *Nat* Neurosci, 2018).

**Table S1:** Number of animals used in this study.

| Experiment | Figures | Sample size | Number of groups | Sample number | Sex | Age (Months) | Number of animals |
| --- | --- | --- | --- | --- | --- | --- | --- |
| Figure 1 | Figure 1A | n = 5-9 mice | 6 | 38 | 10 male, 28 female | 2-4 | 38 WT mice |
|  | Figure 1B | n = 10-12 mice | 2 | 22 | 9 male, 13 female | 2-4 | 22 WT mice |
|  | Figure 1C | n = 5-9 mice | 3(WT mice are same mice from Figure 1A) | 22 | 12 male, 10 female | 2-4 | 14 WT mice, 8 *Vgat-Cre; Oprm1<sup>fl/fl</sup>* mice |
|  | Figure 1D | n = 9 mice | 2(WT mice are same mice from Figure 1A) | 18 | 14 male, 4 female | 2-4 | 9 WT mice, 9 *Sst-Cre; Oprm1<sup>fl/fl</sup>* mice |
|  | Figure 1E | n = 5 mice | 5 | 25 | 18 male, 7 female | 2-4 | 25 WT mice |
|  | Figure 1F | n = 6-7 mice | 2 | 13 | 8 male, 5 female | 2-4 | 6 WT mice, 7 *Vgat-Cre; Oprm1<sup>fl/fl</sup>* mice |
| Figure 2 | Figure 2A | n = 7-9 mice | 2 | 16 | 5 male, 9 female | 2-4 | 16 WT mice |
|  | Figure 2B | n = 6-8 mice | 2 | 14 | 8 male, 6 female | 2-4 | 14 WT mice |
|  | Figure 2C | n = 10-13 mice | 2 | 23 | 10 male, 13 female | 2-4 | 10 WT mice, 13 *Trpv1-Cre; Oprm1<sup>fl/fl</sup>* mice |
|  | Figure 2D | n = 10 mice | 2 | 20 | 15 male, 5 female | 2-4 | 20 WT mice |
|  | Figure 2E | n = 5 mice | 2 | 10 | 7 male, 3 female | 2-4 | 5 WT mice, 5 *Vgat-Cre; Oprm1<sup>fl/fl</sup>* mice |
| Figure 3 | Figure 3A-C | n = 7-11 mice | 4 | 36 | 22 male, 14 female | 2-4 | 10 WT mice, 7 *Vgat-Cre; Oprm1<sup>fl/fl</sup>* mice, 8 *Sst-Cre; Oprm1<sup>fl/fl</sup>* mice, 11 *Trpv1-Cre; Oprm1<sup>fl/fl</sup>* mice |
| Figure 4 | Figure 4A,B | n = 4 mice | 1 | 4 | 4 male | 2-4 | 4 *Vgat-Cre; tdTomato* mice |
| Figure 5 | Figure 5A | n = 19 neurons from 5 mice | 1 | 19 | 3 male, 2 female | 1-2 | 5 *Vgat-Cre; tdTomato* mice |
|  | Figure 5B,C | n = 15 neurons from 5 mice | 1 | 15 | 3 male, 2 female | 1-2 | 5 *Vgat-Cre; tdTomato* mice |
|  | Figure 5D,E | n = 9-14 neurons from 5-6 mice | 2 | 10,15 | 7 male, 4 female | 1-2 | 11 *Vgat-Cre; Ai32* mice |
| Figure 6 | Figure 6A | n = 6-10 mice | 2 | 16 | 6 male, 10 female | 2-4 | 16 WT mice |
|  | Figure 6B | n = 6-7 mice | 2 | 13 | 9 male, 4 female | 2-4 | 13 WT mice |
|  | Figure 6C | n = 6-7 mice | 2 | 13 | 9 male, 4 female | 2-4 | 13 WT mice |
|  | Figure 6D | n = 7 mice | 2 | 14 | 9 male, 5 female | 2-4 | 7 WT mice, 7 *Vgat-Cre; Oprm1<sup>fl/fl</sup>* mice |
| Figure 7 | Figure 7A,B | n = 8-9 mice | 2 | 17 | 8 male, 9 female | 2-4 | 8 WT mice, 9 *Vgat-Cre; Oprm1<sup>fl/fl</sup>* mice |
|  | Figure 7C | n = 5-6 mice | 2 | 11 | 7 male, 4 female | 2-4 | 5 WT mice, 6 *Vgat-Cre; Oprm1<sup>fl/fl</sup>* mice |
|  | Figure 7D | n = 5-6 mice | 2 | 11 | 7 male, 4 female | 2-4 | 5 WT mice, 6 *Vgat-Cre; Oprm1<sup>fl/fl</sup>* mice |
| Figure 8 | Figure 8A | n = 6-10 mice | 3 | 22 | 4 male, 18 female | 2-4 | 22 WT mice |
|  | Figure 8B | n = 9 mice | 2 | 18 | 6 male, 12 female | 2-4 | 18 WT mice |
|  | Figure 8C | n = 8 mice | 3 | 24 | 12 male, 12 female | 2-4 | 24 NOD. CB-17-Prkdc<sup>scid</sup> mice |
|  | Figure 8D | n = 13-14 mice | 2 | 27 | 13 male, 14 female | 2-4 | 27 NOD. CB-17-Prkdc<sup>scid</sup> mice |
| Figure S1 | Figure S1A,B | n = 7-16 mice | 4 (some mice are same mice from Figure 3) | 46 | 27 male, 19 female | 2-4 | 8 WT mice, 15 *Vgat-Cre; Oprm1<sup>fl/fl</sup>* mice, 16 *Sst-Cre; Oprm1<sup>fl/fl</sup>* mice, 7 *Trpv1-Cre; Oprm1<sup>fl/fl</sup>* mice |
|  | Figure S1C | n = 7-10 mice | 4 (same mice from Figure 3) | 33 | 19 male, 14 female | 2-4 | 10 WT mice, 7 *Vgat-Cre; Oprm1<sup>fl/fl</sup>* mice, 8 *Sst-Cre; Oprm1<sup>fl/fl</sup>* mice, 8 *Trpv1-Cre; Oprm1<sup>fl/fl</sup>* mice |
| Figure S2 | Figure S2 | n = 4 mice | 1 | 4 | 4 male | 2-4 | 4 WT mice |
| Figure S3 | Figure S2 | n = 4 mice | 1 | 4 | 4 male | 2-4 | 4 WT mice |
| Figure S4 | Figure S2 | n = 4 cultures | 1 | 4 | 4 male | 2-4 | 4 WT mice |
| **Total number of mice** |  |  |  |  | 243 male, 228 female |  | 471 mice |

## References

Acton D, Ren X, Di Costanzo S, Dalet A, Bourane S, Bertocchi I, et al. Spinal Neuropeptide Y1 Receptor-Expressing Neurons Form an Essential Excitatory Pathway for Mechanical Itch. Cell Rep 2019; 28(3): 625–39 e6.

Andrade A, Denome S, Jiang YQ, Marangoudakis S, Lipscombe D. Opioid inhibition of N-type Ca2+ channels and spinal analgesia couple to alternative splicing. Nat Neurosci 2010; 13(10): 1249–56.

Bardoni R, Shen KF, Li H, Jeffry J, Barry DM, Comitato A, et al. Pain Inhibits GRPR Neurons via GABAergic Signaling in the Spinal Cord. Sci Rep 2019; 9(1): 15804.

Bourane S, Duan B, Koch SC, Dalet A, Britz O, Garcia-Campmany L, et al. Gate control of mechanical itch by a subpopulation of spinal cord interneurons. Science 2015; 350(6260): 550–4.

Brune A, Metze D, Luger TA, Stander S. [Antipruritic therapy with the oral opioid receptor antagonist naltrexone. Open, non-placebo controlled administration in 133 patients]. Hautarzt 2004; 55(12): 1130–6.

Carstens EE, Carstens MI, Simons CT, Jinks SL. Dorsal horn neurons expressing NK-1 receptors mediate scratching in rats. Neuroreport 2010; 21(4): 303–8.

Cevikbas F, Lerner EA. Physiology and Pathophysiology of Itch. Physiol Rev 2019.

Chamessian A, Young M, Qadri Y, Berta T, Ji RR, Van de Ven T. Transcriptional Profiling of Somatostatin Interneurons in the Spinal Dorsal Horn. Sci Rep 2018; 8(1): 6809.

Chen G, Kim YH, Li H, Luo H, Liu DL, Zhang ZJ, et al. PD-L1 inhibits acute and chronic pain by suppressing nociceptive neuron activity via PD-1. Nat Neurosci 2017; 20(7): 917–26.

Chen G, Park CK, Xie RG, Berta T, Nedergaard M, Ji RR. Connexin-43 induces chemokine release from spinal cord astrocytes to maintain late-phase neuropathic pain in mice. Brain 2014; 137(Pt 8): 2193–209.

Corder G, Castro DC, Bruchas MR, Scherrer G. Endogenous and Exogenous Opioids in Pain. Annu Rev Neurosci 2018; 41: 453–73.

Davidson S, Zhang X, Khasabov SG, Simone DA, Giesler GJ, Jr. Relief of itch by scratching: state-dependent inhibition of primate spinothalamic tract neurons. Nat Neurosci 2009; 12(5): 544–6.

Duan B, Cheng L, Ma Q. Spinal Circuits Transmitting Mechanical Pain and Itch. Neurosci Bull 2018; 34(1): 186–93.

Ekelem C, Juhasz M, Khera P, Mesinkovska NA. Utility of Naltrexone Treatment for Chronic Inflammatory Dermatologic Conditions: A Systematic Review. JAMA Dermatol 2019; 155(2): 229–36.

Farmer WS, Marathe KS. Atopic Dermatitis: Managing the Itch. Adv Exp Med Biol 2017; 1027: 161–77.

Gao T, Ma H, Xu B, Bergman J, Larhammar D, Lagerstrom MC. The Neuropeptide Y System Regulates Both Mechanical and Histaminergic Itch. J Invest Dermatol 2018; 138(11): 2405–11.

Han L, Ma C, Liu Q, Weng HJ, Cui Y, Tang Z, et al. A subpopulation of nociceptors specifically linked to itch. Nat Neurosci 2013; 16(2): 174–82.

Han Q, Liu D, Convertino M, Wang Z, Jiang C, Kim YH, et al. miRNA-711 Binds and Activates TRPA1 Extracellularly to Evoke Acute and Chronic Pruritus. Neuron 2018; 99(3): 449–63 e6.

Haring M, Zeisel A, Hochgerner H, Rinwa P, Jakobsson JET, Lonnerberg P, et al. Neuronal atlas of the dorsal horn defines its architecture and links sensory input to transcriptional cell types. Nat Neurosci 2018; 21(6): 869–80.

Huang J, Polgar E, Solinski HJ, Mishra SK, Tseng PY, Iwagaki N, et al. Circuit dissection of the role of somatostatin in itch and pain. Nat Neurosci 2018; 21(5): 707–16.

Ikoma A, Steinhoff M, Stander S, Yosipovitch G, Schmelz M. The neurobiology of itch. Nat Rev Neurosci 2006; 7(7): 535–47.

Ji RR. Recent Progress in Understanding the Mechanisms of Pain and Itch: the Second Special Issue. Neurosci Bull 2018; 34(1): 1–3.

Ji RR, Zhang Q, Law PY, Low HH, Elde R, Hokfelt T. Expression of mu-, delta-, and kappa-opioid receptor-like immunoreactivities in rat dorsal root ganglia after carrageenan-induced inflammation. J Neurosci 1995; 15(12): 8156–66.

Jiang CY, Fujita T, Kumamoto E. Synaptic modulation and inward current produced by oxytocin in substantia gelatinosa neurons of adult rat spinal cord slices. J Neurophysiol 2014; 111(5): 991–1007.

Kardon AP, Polgar E, Hachisuka J, Snyder LM, Cameron D, Savage S, et al. Dynorphin Acts as a Neuromodulator to Inhibit Itch in the Dorsal Horn of the Spinal Cord. Neuron 2014; 82(3): 573–86.

Ko MC, Husbands SM. Effects of atypical kappa-opioid receptor agonists on intrathecal morphine-induced itch and analgesia in primates. J Pharmacol Exp Ther 2009; 328(1): 193–200.

Ko MC, Song MS, Edwards T, Lee H, Naughton NN. The role of central mu opioid receptors in opioid-induced itch in primates. J Pharmacol Exp Ther 2004; 310(1): 169–76.

Kremer AE. What are new treatment concepts in systemic itch? Exp Dermatol 2019.

Kumar K, Singh SI. Neuraxial opioid-induced pruritus: An update. J Anaesthesiol Clin Pharmacol 2013; 29(3): 303–7.

Kwatra SG, Stander S, Kang H. PD-1 Blockade-Induced Pruritus Treated with a Mu-Opioid Receptor Antagonist. N Engl J Med 2018; 379(16): 1578–9.

LaMotte RH, Dong X, Ringkamp M. Sensory neurons and circuits mediating itch. Nat Rev Neurosci 2014; 15(1): 19–31.

Lee H, Ko MC. Distinct functions of opioid-related peptides and gastrin-releasing peptide in regulating itch and pain in the spinal cord of primates. Sci Rep 2015; 5: 11676.

Legroux-Crespel E, Cledes J, Misery L. A comparative study on the effects of naltrexone and loratadine on uremic pruritus. Dermatology 2004; 208(4): 326–30.

Liu Q, Tang Z, Surdenikova L, Kim S, Patel KN, Kim A, et al. Sensory neuron-specific GPCR Mrgprs are itch receptors mediating chloroquine-induced pruritus. Cell 2009; 139(7): 1353–65.

Liu T, Han Q, Chen G, Huang Y, Zhao LX, Berta T, et al. Toll-like receptor 4 contributes to chronic itch, alloknesis, and spinal astrocyte activation in male mice. Pain 2016; 157(4): 806–17.

Liu XY, Liu ZC, Sun YG, Ross M, Kim S, Tsai FF, et al. Unidirectional cross-activation of GRPR by MOR1D uncouples itch and analgesia induced by opioids. Cell 2011; 147(2): 447–58.

Matthes HW, Maldonado R, Simonin F, Valverde O, Slowe S, Kitchen I, et al. Loss of morphine-induced analgesia, reward effect and withdrawal symptoms in mice lacking the mu-opioid-receptor gene. Nature 1996; 383(6603): 819–23.

Metze D, Reimann S, Beissert S, Luger T. Efficacy and safety of naltrexone, an oral opiate receptor antagonist, in the treatment of pruritus in internal and dermatological diseases. J Am Acad Dermatol 1999; 41(4): 533–9.

Mishra SK, Hoon MA. The cells and circuitry for itch responses in mice. Science 2013; 340(6135): 968–71.

Monroe EW. Efficacy and safety of nalmefene in patients with severe pruritus caused by chronic urticaria and atopic dermatitis. J Am Acad Dermatol 1989; 21(1): 135–6.

Moser HR, Giesler GJ, Jr. Itch and analgesia resulting from intrathecal application of morphine: contrasting effects on different populations of trigeminothalamic tract neurons. J Neurosci 2013; 33(14): 6093–101.

North RA, Williams JT. On the potassium conductance increased by opioids in rat locus coeruleus neurones. J Physiol 1985; 364: 265–80.

Pan H, Fatima M, Li A, Lee H, Cai W, Horwitz L, et al. Identification of a Spinal Circuit for Mechanical and Persistent Spontaneous Itch. Neuron 2019; 103(6): 1135–49 e6.

Pavlis J, Yosipovitch G. Management of Itch in Atopic Dermatitis. Am J Clin Dermatol 2018; 19(3): 319–32.

Peer G, Kivity S, Agami O, Fireman E, Silverberg D, Blum M, et al. Randomised crossover trial of naltrexone in uraemic pruritus. Lancet 1996; 348(9041): 1552–4.

Qu L, Fan N, Ma C, Wang T, Han L, Fu K, et al. Enhanced excitability of MRGPRA3- and MRGPRD-positive nociceptors in a model of inflammatory itch and pain. Brain 2014; 137(Pt 4): 1039–50.

Reich A, Szepietowski JC. Opioid-induced pruritus: an update. Clin Exp Dermatol 2010; 35(1): 2–6.

Reszke R, Szepietowski JC. End-Stage Renal Disease Chronic Itch and Its Management. Dermatol Clin 2018; 36(3): 277–92.

Ross SE, Mardinly AR, McCord AE, Zurawski J, Cohen S, Jung C, et al. Loss of inhibitory interneurons in the dorsal spinal cord and elevated itch in Bhlhb5 mutant mice. Neuron 2010; 65(6): 886–98.

Serrano L, Martinez-Escala ME, Zhou XA, Guitart J. Pruritus in Cutaneous T-Cell Lymphoma and Its Management. Dermatol Clin 2018; 36(3): 245–58.

Singh R, Patel P, Thakker M, Sharma P, Barnes M, Montana S. Naloxone and Maintenance Naltrexone as Novel and Effective Therapies for Immunotherapy-Induced Pruritus: A Case Report and Brief Literature Review. J Oncol Pract 2019; 15(6): 347–8.

Sun YG, Chen ZF. A gastrin-releasing peptide receptor mediates the itch sensation in the spinal cord. Nature 2007; 448(7154): 700–3.

Sun YG, Zhao ZQ, Meng XL, Yin J, Liu XY, Chen ZF. Cellular basis of itch sensation. Science 2009; 325(5947): 1531–4.

Tejada MA, Montilla-Garcia A, Cronin SJ, Cikes D, Sanchez-Fernandez C, Gonzalez-Cano R, et al. Sigma-1 receptors control immune-driven peripheral opioid analgesia during inflammation in mice. Proc Natl Acad Sci U S A 2017; 114(31): 8396–401.

Wang Z, Donnelly CR, Ji RR. Scratching after Stroking and Poking: A Spinal Circuit Underlying Mechanical Itch. Neuron 2019; 103(6): 952–4.

Wang Z, Jiang C, He Q, Matsuda M, Han Q, Wang K, et al. Anti-PD-1 treatment impairs opioid antinociception in rodents and nonhuman primates. Sci Transl Med 2020; 12(531).

Xu B, Zhang M, Shi X, Zhang R, Chen D, Chen Y, et al. The multifunctional peptide DN-9 produced peripherally acting antinociception in inflammatory and neuropathic pain via mu- and kappa-opioid receptors. Br J Pharmacol 2020; 177(1): 93–109.

